# CCR2-driven monocyte recruitment is protective against radiotherapy-induced intestinal toxicity

**DOI:** 10.1101/2024.11.05.622080

**Authors:** Nabina Pun, Urszula M Cytlak, Dave Lee, Rita G Domingues, Eleanor J Cheadle, Duncan Forster, Kaye J Williams, Gerard J Graham, Matthew R Hepworth, Mark A Travis, Timothy M Illidge, Douglas P Dyer

**Affiliations:** Wellcome Centre for Cell-Matrix Research, Lydia Becker Institute of Immunology and Inflammation, Faculty of Biology, Medicine and Health, Manchester Academic Health Science Centre, University of Manchester, United Kingdom; Lydia Becker Institute of Immunology and Inflammation, Faculty of Biology, Medicine and Health, Manchester Academic Health Science Centre, University of Manchester, United Kingdom; Targeted Therapy Group, Division of Cancer Sciences, School of Medical Sciences, Faculty of Biology, Medicine and Health, The University of Manchester, Manchester, United Kingdom; Christie NHS Foundation Trust, Manchester Academic Health Science Centre, Manchester, M20 4BX, United Kingdom; Division of Pharmacy and Optometry, Faculty of Biology, Medicine and Health, University of Manchester, Manchester, United Kingdom; Chemokine Research Group, Institute of Infection, Immunity and Inflammation, College of Medical, Veterinary and Life Sciences, University of Glasgow, Glasgow, G12 8TT, United Kingdom; Geoffrey Jefferson Brain Research Centre, Manchester Academic Health Science Centre, Northern Care Alliance NHS Group, University of Manchester, Manchester, United Kingdom

## Abstract

Radiotherapy (RT) is essential in treating abdominal and pelvic cancers but often damages the healthy tissues, particularly the intestines, leading to radiation-induced toxicities with limited treatment options. While the immune system is known to help regulate tissue damage, immune mechanisms involved in RT-induced intestinal toxicity are not fully understood.

Following CT-guided localised intestinal irradiation, single-cell RNA sequencing (scRNA-seq) and flow cytometry revealed RT-induced chemokine-dependent recruitment of innate immune cells. Deletion of C-C chemokine receptor (*Ccr)1*, *Ccr2*, *Ccr3* and *Ccr5*, blocked recruitment and worsened radiation induced toxicities, suggesting an important role for an innate immune cell population in limiting RT-mediated bowel damage.

Furthermore, CCR2-deficient mice showed exacerbated weight loss and intestinal permeability, while the transfer of Ly6C^+^ monocytes alleviated symptoms. Mechanistically, IL-17 cytokine production by group 3 innate lymphoid cells (ILC3s), a critical factor in maintaining intestinal barrier integrity, was found to be reduced in irradiated CCR2^−/−^, however the transfer of Ly6C^+^ monocytes resulted in increased IL-17 levels. These findings demonstrate the critical importance of CCR2-mediated monocyte recruitment in mitigating RT-induced toxicities.

**One Sentence Summary:** CCR2-mediated monocyte recruitment protects against RT-induced intestinal toxicity via IL-17, highlighting a therapeutic target.

## INTRODUCTION

Radiotherapy (RT) is a highly effective non-surgical treatment delivered to over 60% of all cancer patients (*1*), with more than 40% of those cured of their disease having received RT as part of their treatment (*2*). Many of the most common cancers, including prostate, endometrial, cervical, bladder, intestinal, and pancreatic, occur within the abdomen and pelvis and are treated with localised RT as part of the treatment plan (*3*). Despite the effectiveness of RT in local tumour control, the effects on surrounding normal healthy tissues, particularly the intestines can result in severe acute and late toxicity. This radiation-induced injury often leads to severe complications, with acute intestinal toxicity affecting around 90% of patients either during treatment or shortly after (*3–5*).

RT-induced intestinal toxicity is often debilitating, manifesting as symptoms such as diarrhoea, abdominal pain, nausea, and rectal bleeding (*6*, *7*). These symptoms primarily result from damage to the intestinal epithelium, which subsequently compromises the integrity of the intestinal barrier and triggers inflammatory responses (*8*). Radiation-induced damage to healthy intestinal tissue also limits the total therapeutic dose of RT that can be safely administered and may thus compromise local tumour control (*9*). Despite the prevalence of these side-effects, the underlying mechanisms of RT-induced intestinal toxicity remain poorly understood and underexplored. Additionally, there are currently no effective treatments for these complications, representing a significant clinical unmet need.

Investigation of the transcriptional response following RT-induced intestinal injury has revealed an association between immune cell populations, including macrophages and monocytes, with this injury (*10*). However, the role that immune cells play, and the molecular mechanisms and signals governing their recruitment and function during the response of healthy intestinal tissue to localised RT, remains unclear. Chemokines, which are critical regulators of leukocyte recruitment (*11*, *12*), may play a significant role in this process. The role of chemokines in tumour biology is well studied, however the potential role of chemokines in orchestrating RT-induced intestinal tissue toxicity remains ill-defined.

To address this gap, we developed a preclinical murine model of RT-induced intestinal toxicity, with a focus on the role of chemokine-mediated immune cell recruitment. Using this model, we demonstrated that the chemokine-mediated recruitment of specific innate immune cells, protects mice from intestinal injury following RT. Specifically, CCR2^+^ monocytes are critical for protection against radiation-induced intestinal damage, with CCR2-deficient mice displaying increased sensitivity to radiation, reflected by more severe weight loss and compromised intestinal barrier function. Moreover, adoptive transfer of Ly6C^+^ monocytes into CCR2-deficient mice rescues these animals from radiation-induced weight loss and intestinal permeability, underscoring the protective function of this monocyte recruitment.

Our findings establish a direct link between chemokine-mediated monocyte recruitment and protection against RT-induced intestinal injury. Importantly, we also identified a novel mechanism through which CCR2^+^ monocytes exert their protective effect, by promoting IL-17 production by group 3 innate lymphoid cells (ILC3s), which is crucial for maintaining intestinal barrier integrity (*13–17*). These mechanistic insights potentially open new avenues for therapeutic interventions aimed at enhancing CCR2-mediated monocyte recruitment or targeting IL-17 production to alleviate RT-induced intestinal toxicity and improve cancer patient outcomes.

## RESULTS

### Establishment of a localised RT-induced intestinal toxicity model

RT-induced damage to healthy tissues in cancer treatment leads to toxicity and is associated with the activation of an immune response (*4*, *18*). To investigate the role of the immune system in regulating RT-induced toxicity, we developed a mouse model of localised intestinal RT, utilising the small animal radiation research platform (SARRP) to deliver computed tomography (CT)-scan image-guided precise X-ray irradiation. Here, the radiation field was delivered by a 10mm^2^ collimator, ensuring the radiation dose was restricted to the intestines, sparing the bone marrow and other major organs (Fig. S1). This targeted approach to the intestines contrasts with previous studies that have generally utilised total body irradiation alone or that have used limited lead shielding that is much likely to produce RT induced systemic effects distal to the targeted intestines (*19–21*).

To determine the accuracy and localisation of RT dose delivery we used cone beam imaging (Fig. S1) and confirmed the accuracy by analysing the level and localisation of γH2AX staining, a marker of double-stranded DNA breaks (*22–24*), at 6 hours post-RT. Irradiation resulted in significant DNA damage, as evidenced by enhanced γH2AX levels in the small intestines and proximal large intestines (Fig. 1A-C). In contrast, non-irradiated control mice (sham) exhibited minimal DNA damage. γH2AX levels were notably higher in the proximal large intestines, which were within the radiation field, compared to the distal large intestines, which was outside the irradiation field and showed expression levels similar to sham mice (Fig. 1D-E; Fig. S1). These data confirm the precision of our SARRP approach, with minimal scatter radiation outside the targeted area.

**Fig. 1.**
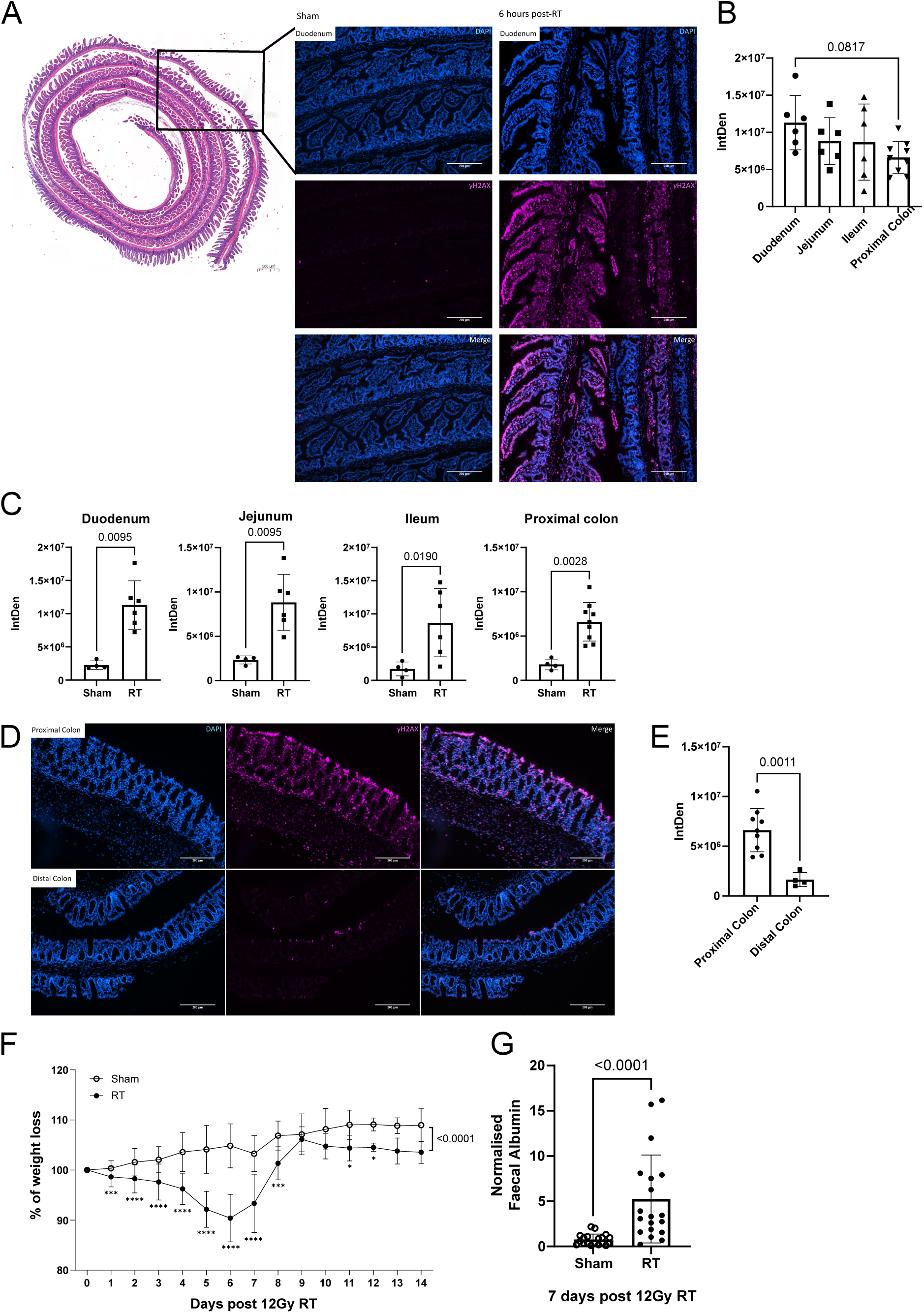
Localised intestinal irradiation results in acute RT-induced toxicity. (A) Left image: Representative H&E-stained ‘swiss-rolled’ duodenum highlighting regions from which snapshots of γH2AX and DAPI stained sections were taken. Right image: Representative of individual DAPI and γH2AX immunohistochemistry stains and a combined merge of the duodenum from sham and irradiated C57BL/6 mice (6 hour post-RT). (B) To assess the accuracy of the SARRP approach, the integrated density (IntDen) of γH2AX was assessed across all intestinal regions in the irradiated mouse. P value was quantified using an ordinary one-way ANOVA, >n = 6 per intestinal region. (C) IntDen of γH2AX in sham vs irradiated mice across each of the four intestinal regions. (D) Confocal microscopy snapshots of DAPI and γH2AX single stain and merge images from the proximal colon (within the irradiated field), and the distal colon (outside of the irradiated field). (E) γH2AX intensity in the proximal colon compared to the distal colon, all in irradiated mice at 6 hours post-RT. (F) Change in body weight of sham and 12 Gy irradiated mice, weights were recorded daily for up to 14 days. (B-E) Data combined from 2 independent experiments, at least n = 4 in RT group and n = 4 sham. (G) Faecal albumin concentrations were normalised to the average of sham controls from the same experiment. Data combined from more than 3 independent experiments, n = 19 irradiated and n = 19 shams. (B-F, H) Data was analysed using the Mann-Whitney t-test. Each data point represents a measurement from a single mouse (G) Data was analysed using a two-way ANOVA with mixed-effects analysis, *P < 0.05, **P < 0.01, ***P < 0.001 and ****P < 0.0001. P value summary = <0.0001 from mixed effects analysis (time and column factor) between RT and sham group. All numerical data represent mean (±SD).

We next assessed whether the localised delivery of RT in our model resulted in tissue damage and associated toxicity. A single 12 Gy fraction of localised RT caused acute body weight loss in mice, peaking at day 6 post-irradiation, with full recovery back to baseline levels by day 8 post-RT (Fig. 1F). To assess the impact of RT on intestinal barrier integrity, we measured faecal albumin levels, a well-established marker of intestinal permeability (*25*). 7 days post-irradiation, faecal albumin concentrations were significantly greater in irradiated mice compared to sham-treated controls (Fig. 1G), indicating that localised RT compromises intestinal barrier function.

We further investigated the effects of radiation on the highly proliferative crypt cells, damage to which has previously been associated with RT-induced intestinal toxicity (*19*, *26*). Sections of the small intestine were stained with haematoxylin and eosin (H&E) and the intestinal epithelium was analysed. H&E staining of small intestine sections revealed significant epithelial damage, with a marked loss of crypts at day 7 post-irradiation (Fig. S2A-B). These findings confirm that our preclinical model successfully replicates RT-induced intestinal toxicity, as evidenced by weight loss (Fig. 1F), increased gut permeability (Fig. 1G), and damage to the intestinal epithelium (Fig. S2A-B) following a 12 Gy dose of abdominal irradiation.

### Local intestinal RT induces an acute innate immune response in the small intestine

The RT-induced host immune response appears to be a key driver of intestinal epithelial injury (*13*, *27*). However, whether the intestinal toxicity observed after localised intestinal RT (Fig. 1F-G, Fig. S2A-B) is immune-mediated, and the immune cells involved, is not known. To address the potential role of the immune response in promoting RT-induced acute intestinal injury, we initially performed unbiased single-cell RNA sequencing (scRNAseq) of CD45^+^ and CD45^−^ cells from the small intestine, at rest and 5 days post-RT. This approach produced a comprehensive cell atlas, encompassing a diverse range of stromal and immune cell populations in both resting and post-RT conditions (Fig. 2A). Examination of cell proportions revealed an expansion of macrophage populations (Fig. S3A), particularly within macrophage clusters 1, 2, and 3 (Fig. 2B). Notably, the macrophage 3 cluster exhibited significant transcriptional changes in response to RT, with numerous genes differentially expressed following irradiation (Fig. 2C), as has been recently observed in radiation-induced salivary gland injury (*28*).

**Fig. 2.**
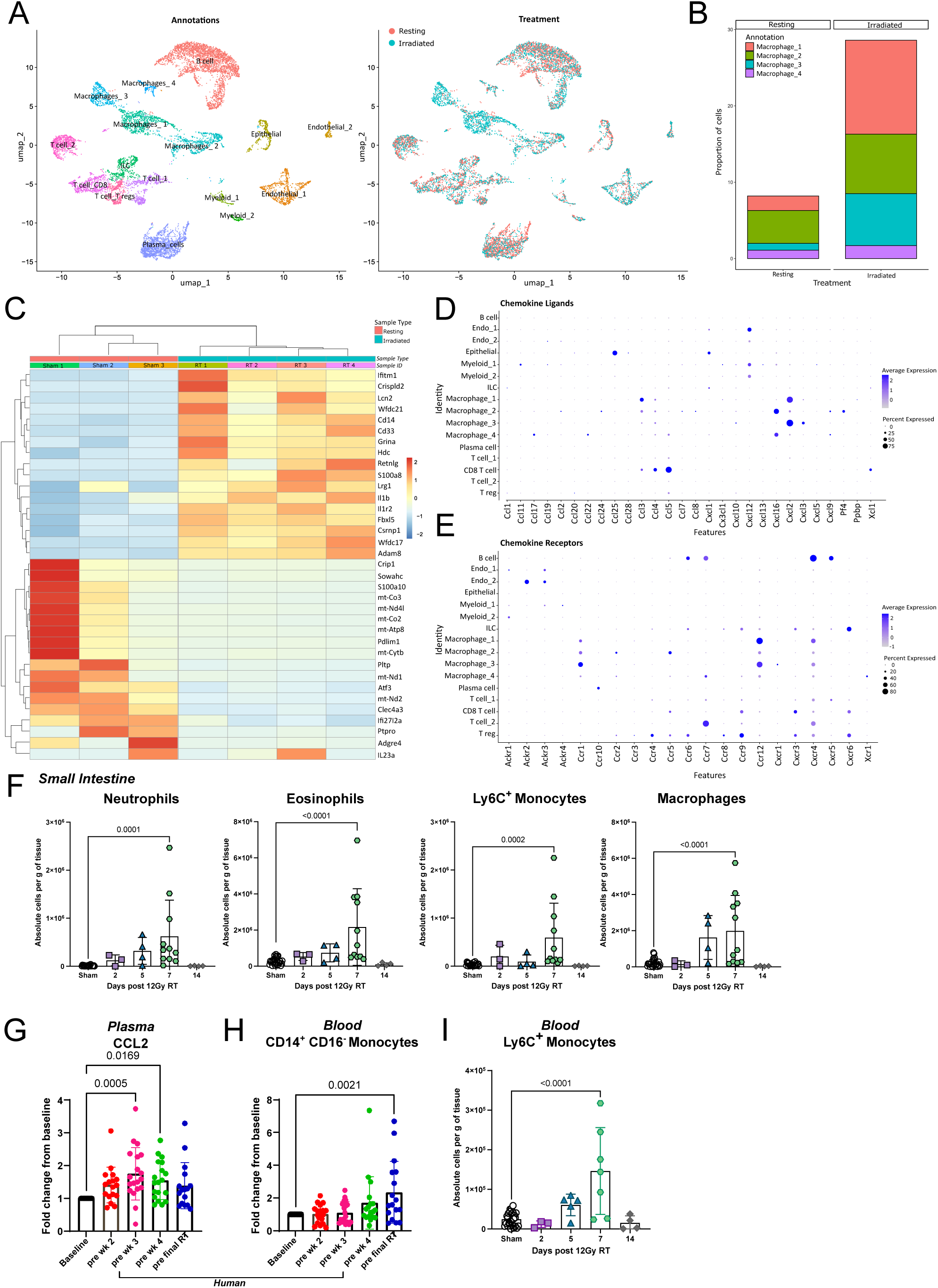
Local intestinal irradiation results in a distinct change in the immune environment within the small intestine. (A) UMAP projection of the small intestine cells profiled. *Left UMAP*, cells coloured by Seurat cluster and annotated with major cell types. *Right UMAP*, cells coloured by group (resting and irradiated). (B) Stacked bar chart show the proportions of each cell subset (macrophage 1, macrophage 2, macrophage 3, and macrophage 4) in resting and irradiated samples. (C) Heatmap displaying the z-score normalised mean expression of the top 40 genes in the macrophage 3 cluster. (A-C) All groups displayed for healthy intestinal samples (resting) and irradiated intestinal samples at day 5 post 12 Gy RT. Bubble plots showing the changes in the proportions of chemokine ligand (D) and receptor (E) in major intestinal cell types at day 5 post 12 Gy. (F) Flow-cytometric analysis of immune cells in the small intestine of sham and irradiated mice at 2-, 5-, 7-, and 14-days post 12 Gy RT. (G) Analysis of human CCL2 levels in plasma and (H) flow cytometric analysis of blood CD14^+^ CD16^−^ monocytes from bladder cancer patients at baseline and before receiving the scheduled fractionated RT dose at pre week (wk) 2, pre wk 3, pre wk 4 and pre final RT. (I) Flow-cytometric analysis of Ly6C^+^ monocytes in the blood of sham and irradiated mice at 2-, 5-, 7-, and 14-days post 12 Gy RT. (F, I) All time points include at least n = 3 irradiated mice for each time point and n = 28 sham mice. (F, G, I) Data was analysed with one-way ANOVA with Dunnett’s multiple comparisons test, with all time points compared to sham.

Specifically, cells in macrophage cluster 3 up-regulated the pro-inflammatory cytokine IL-1β following RT (Fig. 2C) and the highest expression of the neutrophil recruiting pro-inflammatory chemokines *Cxcl2* and *Cxcl3* (Fig. 2D). In contrast, macrophage clusters 1, 2 and 4 displayed no significant changes in gene expression after RT. Interestingly, the macrophage 2 cluster had the highest expression of *Ccr2*, indicating that this population likely contains macrophages recently differentiated from monocytes recruited from the bloodstream (Fig. 2E) (*29*). Gene ontology (GO) term analysis of the different macrophage subtypes highlighted a range of GO terms associated with these populations (Fig. S3B-E). In particular, terms associated with immune cell migration, function and chemotaxis. This is supported by the expression analysis of chemokine ligands (Fig. 2D) and receptors (Fig. 2E) by macrophages and across multiple cell types in the atlas post-RT.

Given the changes observed in innate immune cells from scRNA-seq, we used flow cytometry to fully characterise the numbers and phenotypes of innate immune cells over time, at 2, 5-, 7-, 14-, and 29-days post-RT (gating strategy in Fig. S4). Crucially this approach enabled analysis of cells that are not always captured by scRNAseq, such as neutrophils and eosinophils. Flow cytometry revealed significant increases in the levels of eosinophils, monocytes, macrophages, and neutrophils within the small intestine at day 7 post-RT, with levels returning to baseline by day 14 post-RT (Fig. 2F).

To assess the potential clinical relevance of these immune changes, we analysed peripheral blood mononuclear cells (PBMCs) and plasma from bladder cancer patients treated with abdomino-pelvic RT (gating strategy in Fig. S5). Human plasma analysis revealed elevated levels of the monocyte recruiting chemokine CCL2 (Fig. 2G). Furthermore, flow cytometric analysis of human PBMCs demonstrated increased levels of monocytes (CD14^+^CD16^−^ (*30*, *31*)) in bladder cancer patients treated with RT (Fig. 2H). This observation was mimicked in our preclinical model with elevated blood monocytes at day 7 post-RT (Fig. 2I). Together, the elevated circulating monocyte and CCL2 levels observed in these patients are indicative that RT induces innate immune cell mobilisation from the bone marrow and recruitment to tissues. This data confirms that RT leads to immune cell recruitment in preclinical models and likely also in human patients.

### The inability to recruit innate immune cells increases susceptibility to RT-induced intestinal toxicities

Having observed chemokine production and immune cell recruitment in irradiated tissues, we interrogated the importance of chemokine-driven innate immune cell recruitment in modulating RT-induced intestinal toxicity. We hypothesised that the accumulation of innate immune cells would promote or exacerbate RT-induced intestinal damage. To test this, we used a mouse model deficient in the inflammatory chemokine receptor (iCCR) cluster, which includes the genes encoding *Ccr1*, *Ccr2*, *Ccr3*, and *Ccr5* (*29*, *32*) (Fig. 3A). We have previously established that these receptors control the recruitment of several innate immune cells including monocytes and eosinophils, but not neutrophils (*29*, *32*). Flow cytometric analysis of the small intestine revealed that a lack of iCCRs results in an inability to recruit Ly6C^+^ monocytes (Fig. 3B), eosinophils (Fig. 3C), and a reduction in the number of macrophages (Fig. 3D) at acute time points post-RT with no effect on the level of neutrophils (Fig. 3E).

**Fig. 3.**
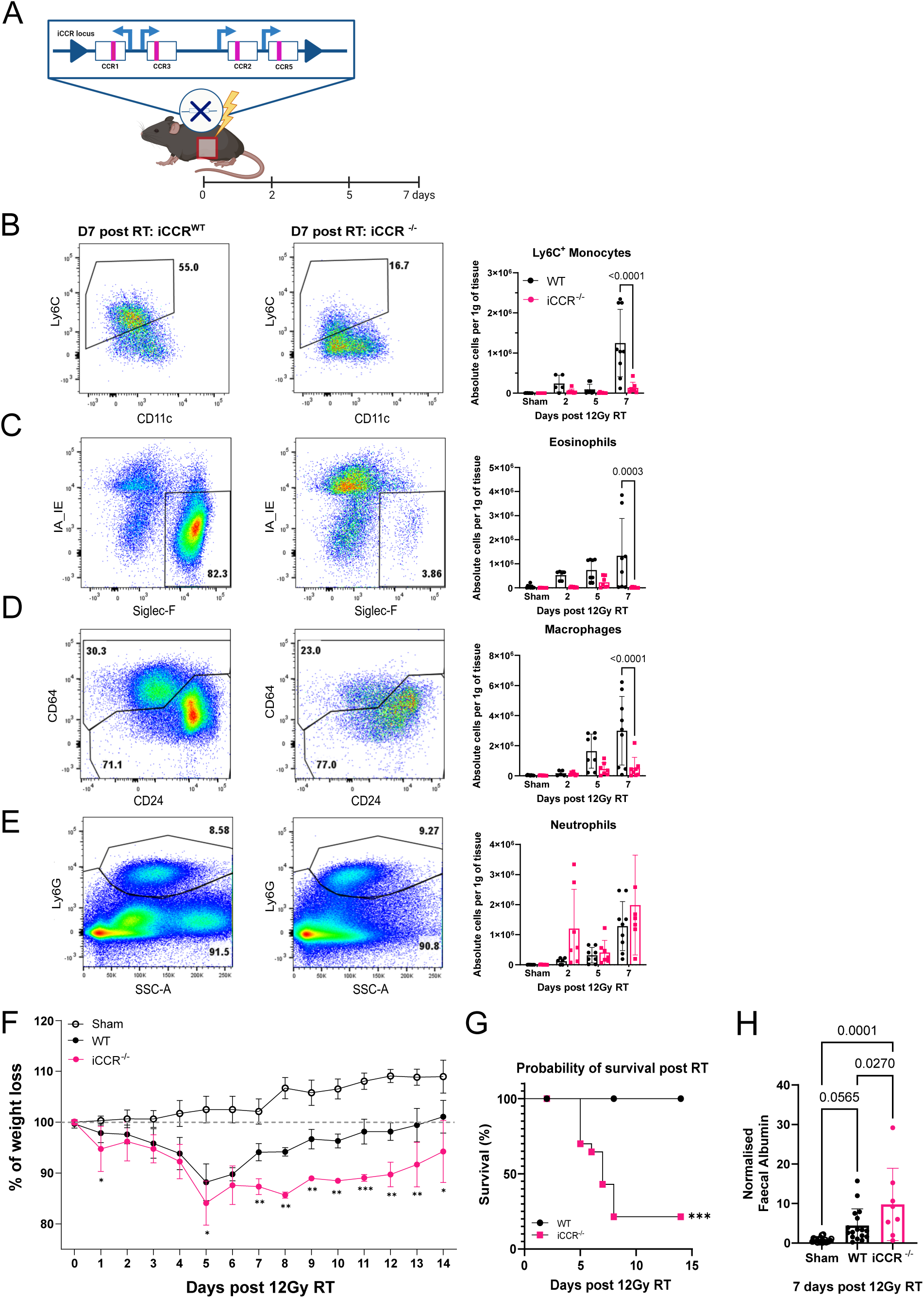
Combined knockout of *Ccr1*, *2*, *3* and *5* results in increased susceptibility to RT-induced toxicity. (A) Localised abdominal X-ray radiation of iCCR^−/−^mice. Mice were culled and small intestine harvested at days 2-, 5-, and 7days post-RT. Representative flow cytometry plots and quantification of (B) Ly6C^+^ monocytes, (C) eosinophils, (D) macrophages, (E) neutrophils in WT and iCCR^−/−^ mice at 2-, 5-, and 7 days post 12 Gy RT. (F) Change in body weight of irradiated WT and iCCR^−/−^ mice following 12 Gy RT in comparison to sham mice, recorded daily for up to 14 days. (G) Percentage of survival following RT over 14 days between irradiated WT and iCCR^−/−^ group. Survival is determined as not losing more than 20% weight which is the humane endpoint in a moderate home office animal license. (H) Faecal albumin concentrations were normalised to the average of sham controls from the same experiment. Normalised faecal albumin levels compared at day 7 post 12 Gy RT across sham, irradiated WT and irradiated iCCR^−/−^. (B-G) Data combined from more than 3 independent experiments, n = 17 irradiated WT, and n = 26. irradiated iCCR^−/−^. (B-E) Data was analysed with two-way ANOVA with Šídák’s multiple comparisons test. (F) P value calculated using a two-way ANOVA with mixed-effects analysis, *P < 0.05, **P < 0.01, and ***P < 0.001. P value summary = <0.0001 from fixed effects analysis (time and column factor). (G) Statistical analysis was carried out using simple survival analysis (Kaplan-Meier) with a log-rank (Mantel-Cox test) curve comparison. (H) Data combined from more than 3 independent experiments, n = 17 irradiated WT, n = 8 irradiated iCCR^−/−^ and n = 19 sham. Statistical analysis using ordinary one-way ANOVA with Tukey’s multiple comparisons test. (B-H) Each data point represents a measurement from a single mouse, and all numerical data represent mean (±SD). Black symbols and bars represent iCCR WT and pink symbols and bars represent iCCR^−/−^.

Contrary to our hypothesis that enhanced immune cell recruitment would promote RT-induced toxicity, iCCR^−/−^ mice, which had reduced innate cell recruitment to the intestine, were more susceptible to RT-induced toxicities, experiencing greater weight loss than the irradiated littermate controls (Fig. 3F), with ∼80% of iCCR^−/−^ mice reaching their humane endpoint by 7 days post-RT (Fig. 3G). Furthermore, irradiated mice deficient in the iCCRs displayed higher faecal albumin levels than the irradiated wild-type (WT) controls (Fig. 3H), indicating increased intestinal permeability. These findings suggest that iCCR-driven innate immune cell recruitment plays a protective role against RT-induced intestinal toxicities.

### Localised RT induces changes in immune cell iCCR expression

Having demonstrated the collective importance of iCCR (CCR1, 2, 3 and 5) receptors in RT-induced toxicity, we next aimed to determine which specific iCCR receptors contribute to this protective function. Firstly, we analysed iCCR expression using the newly developed iCCR reporter mouse (Fig. 4A) (*29*). In agreement with our previous data (Fig. 2F), irradiation of these mice induced significant increases in eosinophil, neutrophil, monocyte, and macrophage levels in the small intestine at 7 days post 12 Gy RT (Fig. S6). Notably, a substantial increase in the proportion of Ly6C^+^ monocytes expressing CCR2 (Fig. 4B-C), eosinophils expressing CCR3 (Fig. 4D-E) and macrophages positive for CCR2 was observed (Fig. 4F-G), at day 7 post-RT. In contrast, there was no significant upregulation of CCR1, 2, 3, or 5 expression post-RT by neutrophils (Fig. 4H-I), suggesting that neutrophil recruitment is likely mediated via other known chemokine receptors and/or non-chemokine mediators (*33–35*). Together with the iCCR^−/−^ data (Fig. 3), these data suggest that chemokine-dependent recruitment of monocyte and/or eosinophil may be protective following local intestinal RT.

**Fig. 4.**
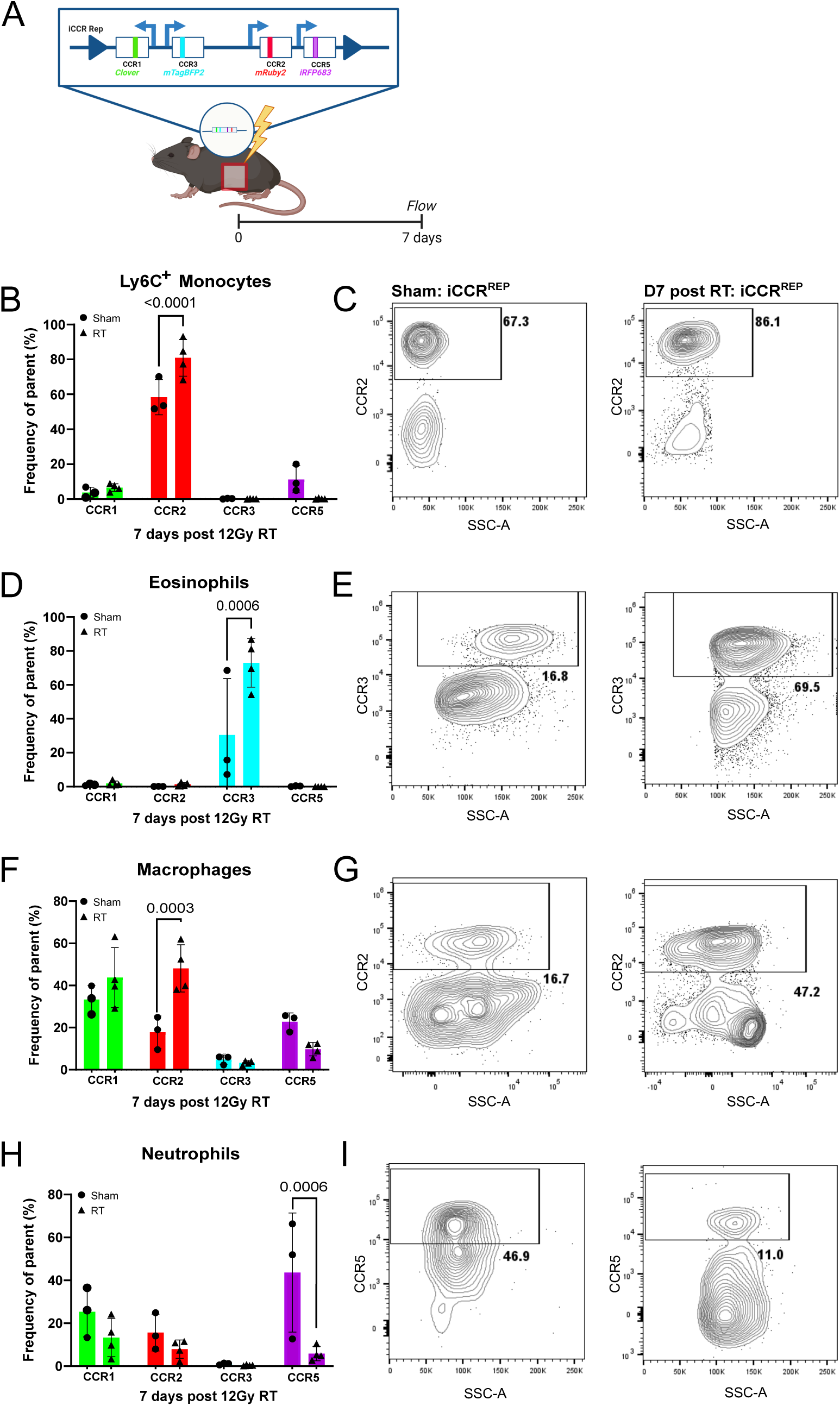
RT induces increased immune cell expression of specific chemokine receptors in the small intestine. (A) iCCR reporters were irradiated with 12 Gy RT and the small intestine was harvested for flow cytometry at day 7 post-RT. Flow-cytometric percentages of (B) Ly6C^+^ monocytes, (D) eosinophils, (F) macrophages and (H) neutrophils expressing CCR1, CCR2, CCR3 and CCR5 in sham and irradiated iCCR reporters at day 7 post-RT. Representative flow cytometry plots of CCR2 expression on (C) Ly6C^+^ monocytes and (G) macrophages in sham and irradiated iCCR reporter mice. (E) Representative flow cytometry plots of CCR3 expression on eosinophils in sham and irradiated iCCR reporter mice. (I) Representative flow cytometry plots of CCR5 expression on neutrophils in sham and irradiated iCCR reporter mice. Data was combined from two independent experiments, n = 3 sham and n = 4 RT iCCR reporters. Data was analysed with two-way ANOVA with Šídák’s multiple comparisons test. Each data point represents a measurement from a single mouse, and all numerical data represent mean (± SD).

### CCR2-driven monocyte recruitment is protective against RT-induced toxicities

With recent literature suggesting potential protective roles of CCR2^+^ Ly6C^+^ monocytes during inflammation (*36*, *37*), we examined the importance and function of monocytes, in the context of RT. To this end, we utilised CCR2^−/−^ mice which have impaired monocyte egress from bone marrow, resulting in reduced monocytes and macrophage populations in the blood and intestines (*38*, *39*). As expected, CCR2 deficiency severely impaired monocyte recruitment after RT (Fig. 5A-B), and in turn, lead to a reduced macrophage population in comparison to irradiated WT mice (Fig. 5C-D), whereas neutrophil (Fig. 5E) and eosinophil (Fig. 5F) recruitment remained unaffected.

**Fig. 5.**
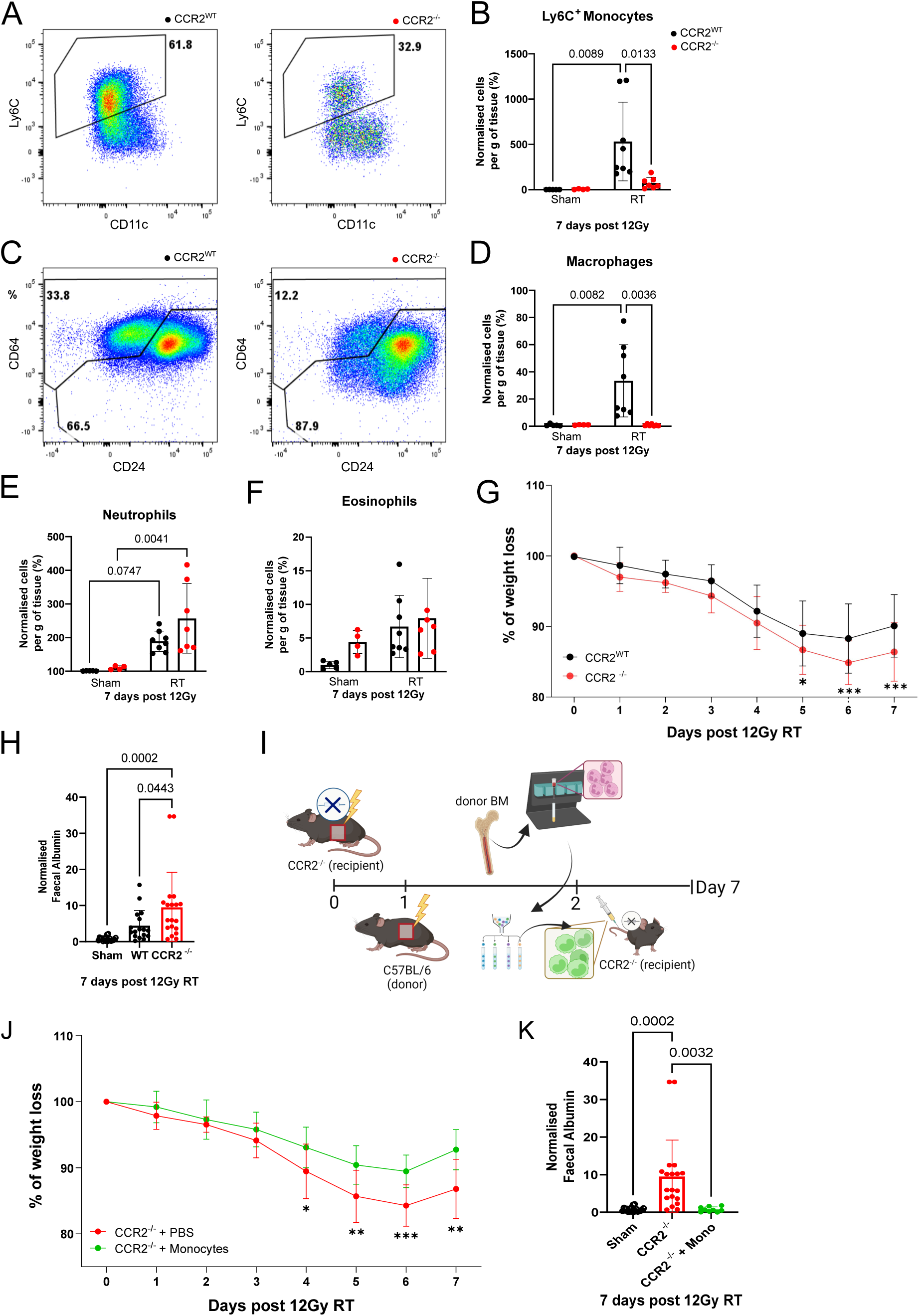
CCR2-recruited monocytes exert protection against RT-induced intestinal toxicity. (A) Representative flow cytometry plots of Ly6C^+^ monocytes in WT and CCR2^−/−^ mice at day 7 post-RT. (B) Normalised (to sham) flow-cytometric quantification of Ly6C^+^ monocytes in WT and CCR2^−/−^ mice at 7 days post 12 Gy RT. (C) Representative flow cytometry plots of CD64^+^ macrophages in WT and CCR2^−/−^ mice at day 7 post-RT. (D) Normalised flow-cytometric quantification of CD64^+^ macrophages in WT and CCR2^−/−^ mice at 7 days post 12 Gy RT. Normalised flow-cytometric quantification of (E) neutrophils and (F) eosinophils in WT and CCR2^−/−^ mice at 7 days post 12 Gy RT. (G) Change in body weight of irradiated littermate WTs and CCR2^−/−^ following 12 Gy RT. Weights were recorded daily for up to 7 days. Data was obtained from more than 3 independent experiments, n = 24 WT, n = 19 CCR2^−/−^. (H) Faecal albumin concentrations were normalised to the average of sham controls from the same experiment. Normalised faecal albumin levels compared at day 7 post 12 Gy RT across sham, irradiated WT and irradiated CCR2^−/−^. Data combined from more than 3 independent experiments, n = 17 irradiated WT, n = 19 irradiated CCR2^−/−^and n = 19 sham. (I) Experimental plan of the adoptive monocyte transfer experiment. (J) Change in body weight of 12 Gy irradiated CCR2^−/−^ mice injected with PBS and CCR2^−/−^ injected with 1×10^6^ Ly6C^+^ monocytes. Weights were recorded daily for up to 7 days. (K) Normalised faecal albumin levels compared at day 7 post 12 Gy RT in sham, irradiated CCR2^−/−^ injected with PBS and irradiated CCR2^−/−^ injected with 1×10^6^ Ly6C^+^ monocytes. (A-F) Data was combined from more than 2 independent experiments and included at least n = 4 in each group. (I-K) Data combined from more than 2 independent experiments, n = 13 sham, n = 14 irradiated CCR2^−/−^ injected with PBS, n = 8 irradiated CCR2^−/−^ injected with 1×10^6^ Ly6C^+^ monocytes. (B, E, D, F, H, K) Statistical analysis using ordinary one-way ANOVA with Tukey’s multiple comparisons test. (G, J) P value calculated using a two-way ANOVA with Šídák’s multiple comparisons test. *P < 0.05, **P < 0.01, and ***P < 0.001. Each data point represents a measurement from a single mouse, and all numerical data represent mean (±SD).

Strikingly, CCR2^−/−^ mice displayed more pronounced weight loss compared to irradiated WT controls, particularly between day 5 to day 7 post-RT (Fig. 5G), akin to iCCR^−/−^ mice. Additionally, CCR2^−/−^ mice showed exacerbated faecal albumin levels, indicating increased intestinal permeability, similar to iCCR^−/−^ mice (Fig. 5H). Collectively, these data suggest that CCR2^+^ monocytes play a crucial role in limiting intestinal toxicity following RT.

### Adoptive monocyte transfer rescues RT-induced intestinal toxicities

To directly test whether CCR2^+^ monocytes have a protective effect in limiting RT-induced gut toxicity, we performed adoptive transfer of monocytes into CCR2^−/−^ mice. Ly6C^+^ monocytes, sorted from irradiated C57BL/6 donor mice (Fig. 5I), were intravenously injected into irradiated monocyte-deficient CCR2^−/−^ recipients 2 days post-RT. We found that transfer of Ly6C^+^ monocytes significantly reduced both the extent of irradiation-induced weight loss (Fig. 5J) and faecal albumin levels (Fig. 5K) in irradiated CCR2^−/−^ mice, compared to PBS-injected controls.

We next tested whether the transfer of Ly6C^+^ monocytes could also protect the iCCR^−/−^ mice, which lacked recruitment of monocytes, eosinophils and exhibited reduced macrophages to the intestine post-RT, from radiation-induced intestinal toxicity (Fig. 2B-D). We found that just the transfer of Ly6C^+^ monocytes into irradiated iCCR^−/−^ mice completely reversed the RT-induced weight loss to that seen in irradiated WT mice (Fig. S7). We also directly tested whether RT-induced iCCR-independent neutrophil recruitment to the intestine could modulate RT-induced intestinal toxicity by depleting neutrophils during RT (Fig. S8A-B). We found that depletion of neutrophils had no effect on RT-induced weight loss (Fig. S8C).

Thus, together, our findings demonstrate that monocytes provide significant protection against toxicities in our pre-clinical model of intestinal RT.

### IL-17 driven regulation of intestinal permeability following RT

We next wanted to determine potential mechanisms via which monocytes exhibit protection from RT-induced damage. Previous literature has suggested that CCR2^+^ monocyte-derived macrophages can regulate intestinal immune responses via effects on ILC3s, including via production of IL-1β (*15*). As described earlier, analysis of our scRNAseq data demonstrated that macrophages in the small intestine upregulated transcription of IL-1β in response to RT (Fig. 2C). In agreement with these results, analysis of small intestine lysates showed significant increases in IL-1β at the protein level following intestinal RT (Fig. 6A). As IL-1β production by macrophages has been shown to regulate IL-17A and IL-22 production by ILC3s (*15*), and these cytokines can play important roles in promoting intestinal repair (*13*, *14*, *16*), we next assessed the production of these molecules by ILC3s in RT-treated WT and CCR2^−/−^ mice by flow cytometry (gating strategy depicted in Fig. S9A). We found that the loss of intestinal barrier integrity in CCR2^−/−^ mice (Fig. 5H) correlated with a reduction in IL-17A (Fig. 6B-C) and IL-22 (Fig. 6B, D) production by CCR6^+^ ILC3s, but not Th17 cells (Fig. S9 B-C), in comparison to irradiated WT control. Crucially, the adoptive transfer of monocytes into mice lacking CCR2 increased IL-17A production (Fig. 6E) but did not alter the potential of CCR6^+^ ILC3s to produce IL-22 (Fig. 6F).

**Fig. 6.**
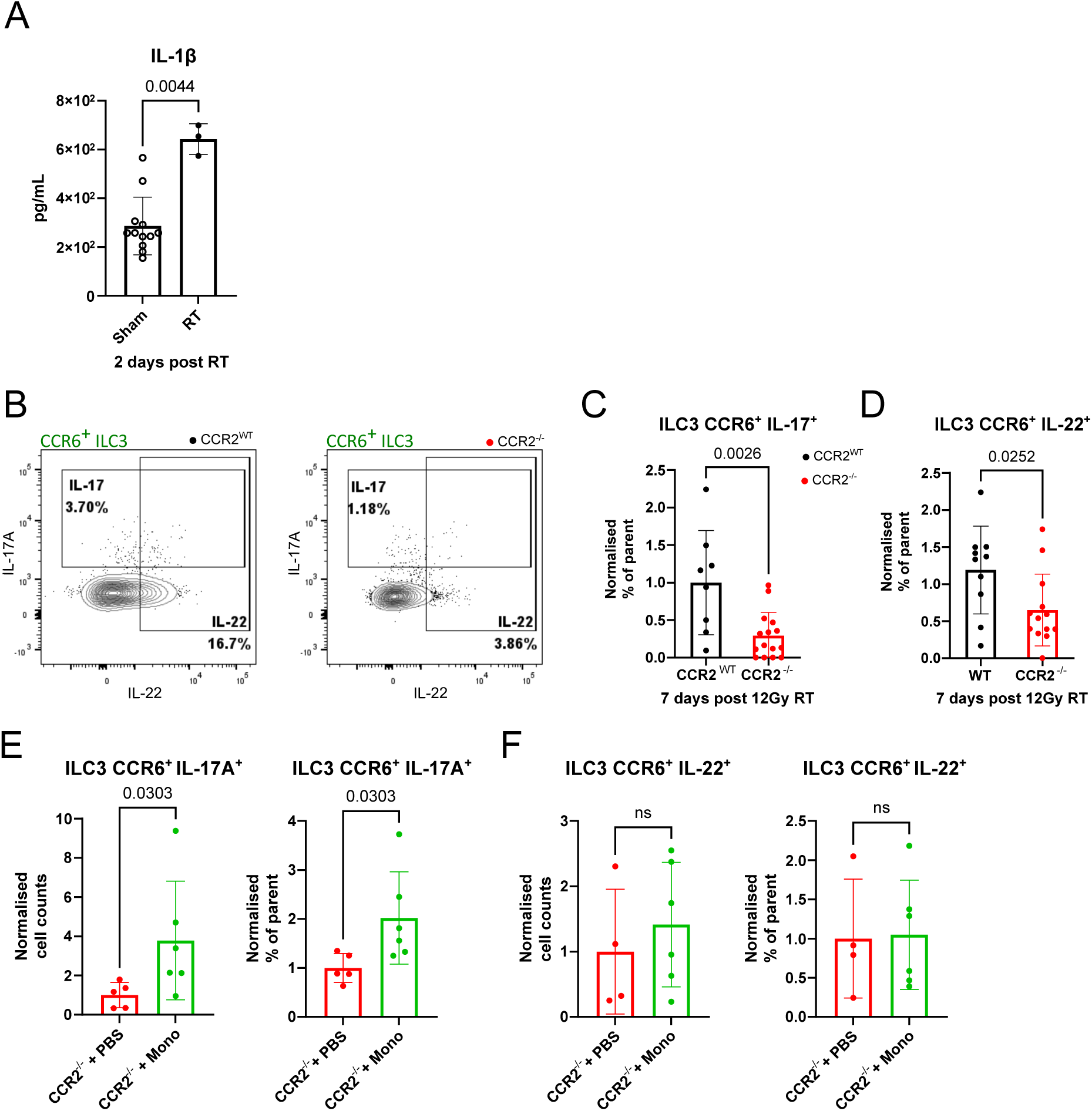
ILC3 activation and IL-17 cytokine production mediates protection following RT-induced intestinal injury. (A) Luminex analysis of IL-1β in the small intestine tissue lysates of sham and irradiated mice at day 2 post 14Gy RT. n = 9 sham and at least n = 3 irradiated mice for each time point. (B) Representative flow cytometry plots of IL-17A and IL-22 expression in CCR6^+^ ILC3. Normalised frequency of ILC3s expressing (C) IL-17A and (D) IL-22. (B-D) Data combined from more than 2 independent experiments and includes at least n = 5 in each group. Normalised cell count and frequency of parent quantification of (E) IL-17 and (F) IL-22 production from CCR6^+^ ILC3 at day 7 post 12 Gy RT. (E-F) All graphed comparisons are between CCR2^−/−^ mice injected with PBS and CCR2^−/−^ mice injected with 1×10^6^ monocytes at 7 days post 12 Gy RT. Data from one experiment, n = 5 for CCR2^−/−^ + PBS, n = 6 for CCR2^−/−^ + Ly6C^+^ monocytes. (A-F) Each data point represents a measurement from a single mouse and all numerical data represent mean (±SD). Data was analysed with an unpaired nonparametric t-test using Mann Whitney test.

Together, these data suggest that monocytes have a critical role in limiting RT-induced intestinal toxicity, which is associated with ILC3-driven IL-17A production.

## DISCUSSION

Earlier diagnosis of cancers has led to an increase in localised treatments including RT resulting in improved cancer survival but associated with an increased incidence and long term morbidity of RT-induced intestinal injury in these cancer survivors (*5–7*). Currently, no effective therapies exist to mitigate the acute and chronic gastrointestinal toxicities associated with RT, underscoring an urgent unmet clinical need to investigate the underlying mechanisms and to develop effective targeted therapeutic interventions.

In this study, we established a localised pre-clinical mouse model of intestinal irradiation that produces hallmark symptoms of RT-induced intestinal toxicity, such as weight loss, increased intestinal permeability, and intestinal epithelial damage. Analysis of this model revealed that innate immune cells rapidly infiltrate the irradiated small intestine, in agreement with previous findings (*10*). In addition, our results demonstrate that monocyte and eosinophil recruitment is driven by CCR1, 2, 3 and 5, as previously suggested in other contexts (*29*, *32*, *40*). Furthermore, we saw no effect of deletion of *Ccr1*, *2*, *3* and *5* on recruitment of neutrophils to the irradiated small intestine despite evidence of their expression of CCR5. These findings advance our understanding of monocyte recruitment and offer novel therapeutic targets to potentially modify RT-induced intestinal inflammation.

Our data demonstrates that CCR2-mediated monocyte recruitment plays a critical role in limiting RT-induced toxicity (weight loss and intestinal barrier integrity). These results are in agreement with emerging evidence highlighting the roles of CCR2^+^ Ly6C^+^ monocytes in various inflammatory processes (*36*, *37*, *41*). However, when considering the therapeutic manipulation of monocyte recruitment to reduce acute and late intestinal toxicity it is also important to consider the potential role of chemokines and the recruited immune cells in the tumour and how this may alter the therapeutic index of RT (*42*). For example, inhibition of the monocyte chemokine receptors CCR2 and 5 has previously been shown to limit radiation-induced monocyte infiltration to pancreatic adenocarcinoma (*43*). This inhibition resulted in improved tumour control and mouse survival when combined with immune checkpoint inhibitors. Indeed, inhibition of CCR2 and monocyte recruitment is generally considered to be beneficial for reducing tumour burden (*44*). Despite the range of preclinical studies suggesting the beneficial outcome of inhibiting CCR2 in different tumour models, human cancer clinical trials failed to produce improved cancer outcomes (*45*). The reasons behind the failure of CCR2 inhibition to influence clinical practice are likely to be complex and include the requirement for ongoing therapeutic intervention with CCR2 inhibition and the inevitable repopulation of these immune subsets once the therapy intervention is stopped. Our study provides additional insights into how potentially CCR2 inhibition may also have adverse effects on healthy tissues, decreasing the therapeutic index by increasing RT-induced toxicity. However, in contrast to ongoing protracted use of CCR2 inhibition for tumour control, promoting monocyte recruitment to decrease radiation induced intestinal injury may only be required for the limited period of the RT delivery. Together these findings suggest that any manipulation of chemokine receptors needs to be carefully considered in enhancing the therapeutic ratio to improve tumour control and decrease normal tissue toxicity.

To understand how monocyte recruitment plays a protective role in response to RT we also investigated potential cellular and molecular mechanisms known to be linked to CCR2^+^ monocytes and regulation of intestinal inflammation. The pro-inflammatory cytokine IL-1β, released by blood monocytes and intestinal macrophages, is crucial for tissue repair and host defence following injury (*46*). We demonstrated an increase in the protein level of this cytokine in the small intestine and increased transcription by an inflammatory macrophage subset in response to RT. IL-1β has previously been shown to stimulate the production of IL-17 and IL-22 by ILC3s, cytokines that regulate intestinal barrier integrity (*13*, *17*, *27*, *47*). We demonstrated that IL-17 and IL-22 secretion by CCR6^+^ ILC3s was regulated by CCR2-mediated monocyte recruitment (Fig. 7). Given the established role for IL-17 and IL-22 in regulating the integrity of the intestinal barrier following infection and injury (*14–17*, *48*, *49*), including in response to RT-induced injury (*50*), we propose that RT-induced infiltrating monocytes play a key role in ILC3-mediated repair of the intestinal epithelium. These findings underscore the importance of Ly6C^+^ monocyte infiltration and differentiation in protecting against radiation-induced intestinal injury. Furthermore, our data also challenge previous suggestions that targeting the IL-17 axis may be useful in improving outcomes to RT (*47*).

**Fig. 7.**
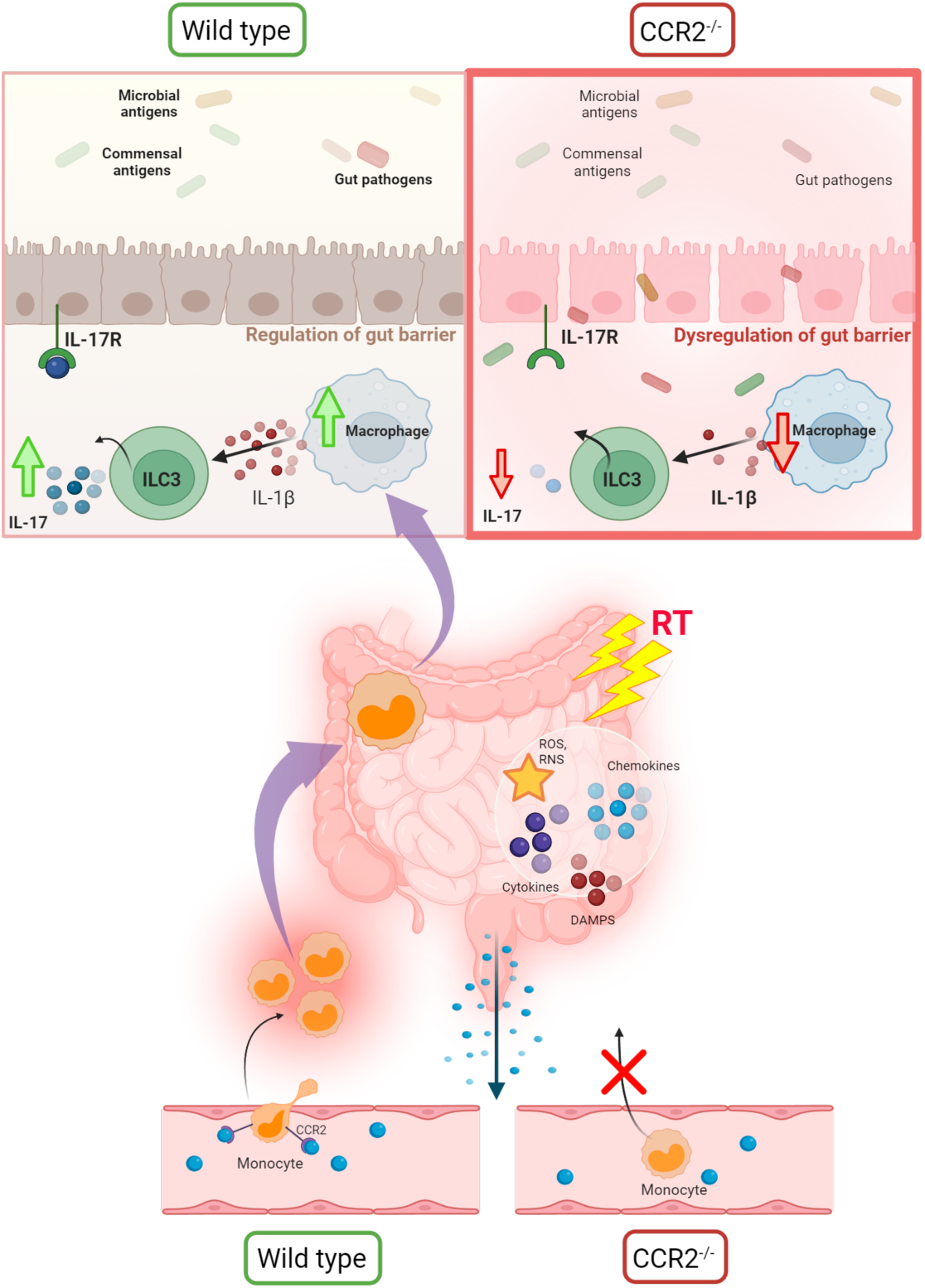
CCR2-mediated monocyte recruitment exerts a protective function following localised intestinal RT. Localised bowel RT results in CCR2-mediated monocyte recruitment into the small intestine. Within the tissues, Ly6C^+^ monocytes differentiate into intestinal macrophages which are known to secrete IL-1β. This increase in IL-1β, production can, in turn, activate ILC3 to produce the cytokines, IL-17 and IL-22, key in regulation of the intestinal barrier. However, CCR2 deficient mice lack the infiltration of Ly6C^+^ monocytes into the small intestine, impacting macrophage numbers. This results in decreased ILC3 production of IL-17, leading to an increase in intestinal barrier permeability. Dysregulation of intestinal epithelial barrier permeability may result in increased translocation of commensal and pathogenic microbes. Figure created with BioRender.com.

In conclusion, through the development of a highly targeted pre-clinical model of RT-induced intestinal toxicity, our findings uncover a critical protective role for Ly6C^+^ monocytes in mitigating radiation-induced intestinal toxicity. The recruitment of these monocytes by CCR2 and their subsequent effect on ILC3s may be key mechanisms underlying this intestinal protection. These results suggest that therapeutic strategies aimed at enhancing CCR2-mediated monocyte recruitment or promoting ILC3 function in non-cancer tissues may represent promising approaches for reducing the major unmet clinical need of RT-induced intestinal injury.

## MATERIALS AND METHODS

### Mice

Up to 5 mice aged between 9 and 16 weeks were housed in individually ventilated cages at the University of Manchester, under specific pathogen-free conditions, standard chow, and water provided ad libitum, constant temperature, and a 12-hour light and dark cycle. Female C57BL/6 mice were purchased from Charles River. iCCR reporter, iCCR^−/−^ and CCR2^−/−^ mice were bred in-house and maintained at the University of Manchester. iCCR^−/−^ and iCCR reporters were previously generated (*29*, *32*). The CCR2^−/−^ strain was a kind gift from Dr Joanne Konkel, University of Manchester. All animal strains were generated and maintained on a C57BL/6 background. For transgenic mouse studies, littermates of the same sex were assigned to experimental groups according to genotype. Where necessary or appropriate both female and male mice were utilised, and sex was not found to be a confounding factor in the observations reported herein. All experiments were carried out following ethical approval from The University of Manchester under a license from the UK Home Office (Scientific Procedures Act 1986).

### Small animal radiation research platform

Anaesthetised mice were transferred to the SARRP (Xstrahl), equipped in nose-cone isoflurane delivery, for CT scan image-guided precise X-ray irradiation. CT scan images were taken from all mice under a constant flow of isoflurane (2%) with oxygen (1L/min) for up to 10 min. ‘Sham’ mice were placed back into their respective cages following CT scan. Single fraction dose (12 Gy) was applied to the abdomen of C57BL/6 or transgenic mice, through a 10 × 10 mm collimator, using a parallel-opposed method delivering a half dose from the left side and a half from the right side of the mouse body. All experimental groups were floor-fed (chow) following RT and monitored daily for changes in weight and appearance.

### Clinical samples

Patients with muscle invasive bladder cancer (T2-T3b, grade 3 receiving 52.5-55Gy RT in 20 fractions plus weekly Gemcitabine (100mg/m^2^)) were recruited to the Manchester Cancer Research Centre Biobank study 18CAWE03 under MCRC Biobank Research Tissue Bank Ethics. Patients were consented in accordance with the declaration of Helsinki. 9ml of blood was collected in a EDTA vacutainer prior to the first dose of RT, then weekly just before the next RT dose, and at the final fraction of RT.

Blood was centrifuged at 2000 x g for 10 min and plasma harvested and stored in 1ml aliquots at −80 °C. The remaining blood was remixed, rocked overnight at RT and then diluted with HBSS (Sigma-Aldirch) to 20 ml before layering over 2 x 4ml lymphoprep (Stem Cell Technologies) and centrifuging at 800 x g for 20 mins. The buffy coat was collected, washed twice in HBSS and frozen at 10^7^ cells/ml in 50% human AB serum (Sigma-Aldrich), 40% RPMI 1640 (Gibco) and 10% dimethyl sulfoxide (DMSO).

### *In vivo* neutrophil depletion

Depletion of neutrophils in female C57BL/6 mice was conducted by intraperitoneal (I.P.) injection with 500 μg anti-Ly6G antibodies or 500 μg IgG isotype control (clone 1A8; BioXCell). I.P. injections of either vehicle or anti-Ly6G mAb started two days prior to single fraction 12 Gy RT and then every two days on alternating sides, no anaesthesia, no analgesia.

### Small intestine immune cell isolation

Cells were isolated as previously described (*51*). Briefly, small, intestines were flushed, and then cut open longitudinally. To remove mucous and epithelial cells, the intestines were washed in 3% RPMI (RPMI supplemented with 10% foetal calf serum (FCS) (Sigma-Aldrich), 1% Penicillin/streptomycin (Sigma-Aldrich), 1% L-Glutamine (Sigma-Aldrich) and 20 mM 4-(2-hydroxyethyl)-1-piperazineethanesulfonic acid (HEPES)) with 1 mM Dithiothreitol (DTT) (Sigma-Aldrich), and 5 mM EDTA (Sigma-Aldrich). Supernatant filtered, and tissues were thoroughly chopped and incubated in cRPMI (RPMI supplemented with 1% non-essential amino acids (Sigma-Aldrich), 1% sodium pyruvate (Sigma-Aldrich), 1% penicillin-streptomycin, 1% L-Glutamine and 24 mM HEPES) with Liberase TL (Roche) and DNase (DN25, Sigma-Aldrich) for 30 min at 37 °C. The supernatant was filtered through 40 μm mesh filter and single-cell suspensions were acquired after centrifugation.

### *Ex vivo* cell stimulation

To assess ILC3 capacity for IL-17A and IL-22 cytokine production, following immune cell isolation and counting, intestinal cells were resuspended in complete media supplemented with 10% FCS, 1% Penicillin/streptomycin, 1% L-Glutamine and plated at a density of 2-3 × 10^6^ cells per well in a round U-bottom Nunclon Delta treated 96-well plates with lids (Thermo Fisher Scientific). Plate was then centrifuged at 500 x g for 5 min at 4 °C before being stimulated with 100 μl of complete media + 1X eBioscience Cell Stimulation Cocktail (containing phorbol 12-myristate 13-acetate (PMA), ionomycin plus protein transport inhibitors brefeldin A and monensin) (Thermo Fisher Scientific) in complete media for 4 hours at 37 °C, with CO_2_. After incubation, cells were centrifuged at 500 x g for 5 min at 4 °C, ready for antibody staining.

### Adoptive transfer of bone marrow Ly6C^+^ monocytes

C57BL/6 mice (“donor mice”) received a single fraction of 12 Gy RT. 24 hours post-irradiation, the donor mice were euthanised, and the femurs and tibias were harvested under aseptic conditions. The surrounding skin and muscle tissue were removed in a sterile environment, and bone marrow cells were extracted from the bones.

The extracted bone marrow cells were lysed in 1 mL of ACK red blood cell (RBC) lysis buffer (Sigma-Aldrich) for 2 min to remove RBCs. Following lysis, the cells were washed with 1X PBS and centrifuged at 500xg for 5 min at 4°C. To enrich the bone marrow cells, magnetic-activated cell sorting (MACS) was performed using Anti-Ly6G Microbeads Ultrapure (Miltenyi Biotec) according to the manufacturer’s protocol. The isolated bone marrow cells were subsequently stained and sorted on the FACSAria cell sorter (BD Biosciences) to isolate Ly6C^+^ monocytes, as detailed below.

Irradiated iCCR/CCR2^−/−^ and iCCR/CCR2 WT mice (at day 2 post-RT) were randomly assigned to experimental groups. Each group was treated with an intravenous injection of either 1 × 10^6^ purified Ly6C^+^ monocytes in 100 μl of sterile 1X PBS or 100 μl of sterile 1X PBS as a vehicle control. Mice were closely monitored for up to 14 days post-treatment.

### Flow cytometric analysis and cell sorting

For mouse bone marrow cell surface staining, single-cell suspensions were incubated with surface protein antibodies at 4°C for 30 min. The following antibodies were used for mouse cell surface staining prior to sorting monocytes: CD45 (1:100 dilution, 30-F11) CD64 (1:100 dilution, X54-5/7.1), Ly6G (1:100 dilution, 1A8), MHC II (1:100 dilution, M5/114.15.2), CD24 (1:200 dilution, M1/69), Ly6C (1:100, HK1.4), Siglec-F (1:100 dilution, E50-2440), CD11b (1:100 dilution, M1/70), CD11c (1:100 dilution, HL3). To discriminate dead cells, 0.25 µg/ml 4′,6-diamidino-2-phenylindole (DAPI) (Sigma-Aldrich) was added to cell surface-stained bone marrow single-cells prior to cell sorting.

To stain freshly isolated intestinal cells, Zombie Fixable Viability Kit (1:2000 dilution, BioLegend) was first applied for 15 min at 4 °C, washed in PBS, and then cells incubated with Fc block for 10 min at 4 °C (1:100 dilution). Once washed in PBS, a cocktail of fluorochrome-conjugated antibodies was applied for 30 min in the dark at 4 °C. The following antibodies were used for mouse cell surface staining cocktail: Tim-4 (1:200 dilution, 21H12), CD64 (1:350 dilution, X54-5/7.1), CD103 (1:350 dilution, M290), CD45 (1:200 dilution, 30-F11), MHC II (1:500 dilution, M5/114.15.2), CD24 (1:200 dilution, M1/69) CD11c (1:350 dilution, HL3), CD4 (1:350 dilution, GK1.5), Ly6C (1:200 dilution, HK1.4), Siglec F (1:350 dilution, E50-2440), γδTCR (1:400 dilution, GL3) F4/80 (1:200 dilution, BM8), CD8a (1:200 dilution, 53-6.7), Ly6G (1:350 dilution, 1A8), CD11b (1:350 dilution, M1/70). Cells were washed and fixed with Foxp3/Transcription Factor Staining Buffer Set (eBioscience) for 20 min or overnight.

For intracellular staining of IL-17, IL-22, RORγt and IFNγ, cells were fixed and permeabilised using Foxp3/Transcription Factor Staining Buffer Set (eBioscience), then stained with the following antibodies: IL-17 (1:100 dilution), IL-22 (1:100 dilution), RORγt (1:200 dilution), IFNγ (1:100 dilution). Cells were analysed by a Symphony flow cytometer (BD Biosciences) with BD FACS DIVA software (v.8.0.2) (BD Biosciences) Flow cytometric data were analysed with FlowJo v.10 software (BD Biosciences).

### Clinical PBMC flow cytometry staining

PBMC were thawed rapidly in a 37°C water bath, washed in RPMI 1640 plus 2mM L-Glutamine, 10% FCS, 50U/ml Benzonase Nuclease (BN) (Sigma-Aldrich) (Complete Media-CM) and then incubated at 37°C in 2 ml CM for one hour. PBMC were then washed, counted and 4×10^5^ were taken forward for flow cytometry. PBMC were washed in PBS plus BN, incubated for 30 mins at room temperature with 1:1000 LIVE/DEAD™ Fixable Blue Dead Cell Stain Kit (Invitrogen) in PBS plus BN, washed in FACS buffer (PBS, 1% FCS, 50U/ml BN), incubated with Trustain FcX Fc block (1:100 in FACS buffer) for 15 min at 4 °C, washed in FACS buffer and then incubated with 100µl of the antibody staining cocktail in FACS buffer for 30 mins at 4°C. The following antibodies were added to the antibody staining cocktail: HLA-DR (1:20 dilution, LN3), LAG-3 (1:20 dilution, 7H2C65), PD-1 (1:20 dilution, EH12.2H7), CD4 (1:80 dilution, A161A1), CD8 (1:80 dilution, SK1), CD16 (1:80 dilution, 3G8), CCR7 (1:20 dilution, 2D1), CD14 (1:40 dilution, HCD14), CD19 (1:80 dilution, HIB19), CD45RO (1:80 dilution, UCHL1), CD3 (1:80 dilution, SK7), CD45RA (1:80 dilution, HI100), CD56 (1:80 dilution, HCD56). Cells were washed in FACS buffer and fixed in 1% paraformaldehyde (BioLegend) and then acquired on a BD Fortessa flow cytometer. Flow cytometric data were analysed with FlowJo v.10 software (BD Bioscience). Compensation controls were created using BD anti-mouse I, and BD anti-Rat Ig k/negative control compensation particles sets (BD Biosciences).

### Tissue homogenisation and Luminex

Liquid nitrogen (LN_2_) was poured onto snap-frozen intestinal tissues and crushed with a pestle and mortar. Samples were homogenised and lysed with a Tris-based lysis buffer (Tissue extraction reagent I, Invitrogen) supplemented with 1mM of phenylmethylsulfonyl fluoride (PMSF, Thermo Scientific) and protease inhibitor cocktail (Pierce Protease Inhibitor Mini Tablets, EDTA-free, Thermo Scientific) before use. 10 mL/g of lysis buffer was added, and samples were placed in a rotisserie to spin at 4 °C for 3 hours. Samples were spun in a pre-cooled centrifuge at 10,000 x g for 15 min at 4 °C. Supernatants were then collected and analysed using a Bio-Plex Pro Mouse Chemokine Panel, 31-Plex Assay (Bio-Rad), per manufacturer’s instructions. Data was acquired on a Bio-Plex Manager™ (Software 6.2)

### Human Luminex

Plasma was thawed at room temperature, pelleted and 25 µl supernatant was analysed in duplicate using a ProcartaPlex Human Cytokine & Chemokine Panel 1A, 34-plex (Invitrogen #EPX340-12167-901) according to the manufacturer’s instructions. Samples were acquired on a Luminex 200 (Millipore).

### Enzyme-linked immunosorbent assay (ELISA)

Specific concentrations of faecal albumin were measured by ELISA, using the Mouse Albumin ELISA Kit (Bethyl Laboratories, Inc) in a 96-well high-binding ELISA plate as per the manufacturer’s instructions. Plates were read on a VersaMax Microplate Reader (Marshall Scientific) at 450 nm.

### Histology

Intestinal tissues were preserved in 10% neutral buffered formalin (Sigma-Aldrich) overnight and then transferred into 70% Ethanol (EtOH), ready for overnight processing using the Leica ASP 300 tissue processor (Leica Biosystems). 5μm formalin-fixed paraffin-embedded (FFPE) transverse sections were cut (Leica RM2255; Leica Biosystems) and mounted on Superfrost Plus Adhesion Slides (25×75mm; Thermo Scientific).

### Immunohistochemistry

To examine the extent of RT exposure, expression of γH2AX, a sensitive molecular marker of DNA damage (*24*), expression was assessed using immunofluorescence. Swiss rolled intestinal tissues were collected at 6 hours post-RT, a time point chosen based on studies that reveal maximal γH2AX expression following RT with residual damage to intestinal crypt cells apparent until 24 hours post-RT (*52*). FFPE tissues were de-waxed in xylene 2 x 5 mins and rehydrated in 100%, 90% (v/v), and 70% (v/v) EtOH for 5 min each and then washed for 1 min in dH20. Slides were placed in Tris EDTA buffer for antigen retrieval (10mM Tris base, 1mM EDTA, 0.05% Tween 20, pH 9.0) at 80% microwave power for 3 min and 40% power for 20 min. Slides were brought to room temperature and transferred to 1X PBS 2 x 5 min. Sections were permeabilised in PBS containing 0.5% Triton X-100 for 20 min and washed with 1X PBS 2 x 5 min. Tissues were enclosed using a PAP pen and blocked with 1% bovine serum albumin (BSA), and 10% goat serum in PBST (PBS + 0.1% Tween 20) for 30 mins followed by two 1XPBS washes. Slides were incubated overnight at 4 °C with the primary anti-γ-H2AX antibody (Phospho-Histone H2A.X (Ser139)) (Cell Signalling Technology), diluted 1:250 (v/v) in blocking buffer. After washing in PBST for 2 x 2 mins, stained slides were then washed in PBS for 5 min and incubated for 1 hour at room temperature with secondary Alexa Fluor 647-conjugated goat anti-rabbit IgG (H+L) antibody (ThermoFisher), diluted 1:200 (v/v) in PBS. After washing in PBS for 3 x 5 min, stained slides were incubated for 2 min with DAPI, diluted 1:5000 (v/v) in PBS. Slides were washed in PBS for 5 min and mounted with 1 drop of Prolong Gold Antifade mountant (ThermoFisher) and cover slipped.

### Image analysis

Imaging was performed using the EVOS M5000 microscope (ThermoFisher) with a 10x objective. Quantification of γH2AX expression in the intestine was performed by analysis of confocal images using ImageJ software. The integrated density (IntDen) measurement was taken, and the background levels were considered by averaging IntDen from negative control sections and subtracting the value from the IntDen of each sample.

### scRNA-seq

Male and female CCR2 WT, iCCR WT, CCR2^−/−^ and iCCR^−/−^ mice were irradiated (n = 3 per group), as previously described an euthanised at 5 days post-RT. Sham mice were anaesthetised, and CT scan images were taken (n = 3 per group). The small intestine was processed into single cell suspensions and stained with CD45 anti-mouse antibody (1:200 dilution, 30-F11). Stained cells were sorted on the FACSAria cell sorter (BD Biosciences) to obtain 50% CD45^+^ and 50% CD45^−^ mixed population. Sorting was performed separately for each mouse. The Chromium Next GEM Single Cell Fixed RNA Sample Preparation Kit, 16 rxns (PN-1000414) was used to fix 1.5 × 10^6^ cells per sample, according to the manufacturer’s protocols. To generate Fixed RNA gene expression libraries, fixed single-cell suspensions were mixed with unique probe barcodes, and hybridised overnight (16-24 hour) at 42 °C. Samples were pooled immediately after hybridisation and washed together, then 1.28 × 10^5^ cells were loaded onto a Chromium next GEM Chip Q for GEM generation and run on the Chromium X (10x Genomics).

### scRNA-seq data analysis - pre-processing

All samples were subjected to processing with Seurat (v5.0.3). Quality control and filtering was performed to retain high-quality cells that had less than 10% mitochondrial content, genes between 200 – 3000 and annotated as a “Singlet” following doublet prediction *via* the singleCellTK package (v2.12.2). Successful cells were transformed and normalised individually (SCTransform) followed by integration of all samples and subsequent dimension reduction and clustering. Due to low cell numbers in some samples, the k.weight parameter was reduced to 20; all other parameters remained as default.

### scRNA-seq data analysis - cluster annotation

Cell identities were inferred based on the consensus agreement between reference atlases within the Tabula muris resource. The integrated Seurat object was subjected to projection mapping onto each of the 16 available reference atlases (representing different tissues) available in this resource; those atlases that exceeded a median prediction score threshold of 0.9 were retained and a cell was assigned an identity based on the top-most occurring annotation across the surviving reference atlases. Finally, a cluster was assigned a cell-type identity if over 60% of cells in that cluster shared the same annotation; those clusters that did not meet this criterion were removed from further consideration.

### scRNA-seq data analysis - differential expression analysis

Identification of differentially expressed genes (DEGs) was performed with the FindMarkers or FindAllMarkers function within Seurat. For identifying discriminatory cell markers between-clusters, the FindAllMarkers function was used, and only cluster-enriched genes were retained. To identify DEGs within the same clusters but across experimental conditions; cells were first pseudo-bulked through aggregation of counts with AggregateExpression and conditions compared using FindMarkers() with DESeq2 as the test mode.

### scRNA-seq data analysis - over-representation analysis

Differentially expressed genes were used for over-representation analysis (ORA) of gene ontology terms *via* the ClusterProfiler package (enrichGO). For cluster-specific annotations, DEGs passing a 5% p-adjusted value in conjunction with a Log2 fold-change greater than 0.5 were used. For condition-comparisons; DEGs that passed a 5% p-adjusted threshold were accepted.

### Statistics

All statistical analysis was performed using GraphPad Prism, v10.1.2 (GraphPad Software). Data was analysed to determine normal distribution according to a variety of normality tests including the Anderson-Darling test, D’Agostino and Pearson test, Shapiro-Wilk test, and Kolmogorov-Smirnov test. This outcome determined subsequent analysis. All normally distributed data was analysed using unpaired parametric student’s t-test (comparison of only two groups), a one-way analysis of variance (ANOVA) (comparison of 3 or more groups with one independent variable) with Dunnets multiple comparison test, or a two-way ANOVA (comparison of more than two groups with two independent variables). When not normally distributed, data was analysed using unpaired nonparametric student’s t-test or one-way ANOVA using the Kruskal Wallis test with Dunnett’s multiple comparison test. Results from these tests were reported as significant if the P was ≤ 0.05, with results in graphs shown as mean ± standard deviation (SD). In figures, *P < 0.05, **P < 0.01, ***P < 0.001 and ****P < 0.0001.

## Acknowledgments

The authors acknowledge members of the Dyer lab, the Targeted Therapy Group (including Dr Shuhui Cheng, Phil Jones and Lisa Cooper) and the Radiotherapy Network (RadNet) team for critical discussion and processing and analysis of clinical samples. We acknowledge members of the biological services, flow cytometry, histology and bioimaging facilities at the University of Manchester and CRUK Manchester Institute. We acknowledge the MCRC Biobank, Prof Catharine West and research nurses from the Christie NHS Foundation Trust for assistance with collection of patient samples. Diagrams created using Biorender.com

## Author contributions

Conceptualization: NP, DPD, TMI, MAT, KJW

Methodology: NP, DL, UMC, EJC, RGD, GJG, MRH, DF

Investigation: NP, UMC, EJC, RGD, MRH

Visualization: NP

Funding acquisition: TMI, DPD, MAT, KJW

Project administration: TMI, MAT, DPD

Supervision: TMI, DPD, MAT

Writing – original draft: NP, DPD

Writing – review & editing: NP, DPD, UMC, MAT, TMI

## Data and materials availability

Any data not explicitly evident in the paper will be available on reasonable request.

