## Supplementary information for "CCR2-driven monocyte recruitment is protective against radiotherapy-induced intestinal toxicity"

### SUPPLEMENTARY MATERIALS AND METHODS

#### Haematoxylin and eosin (H&E) staining

Slides were dewaxed in xylene 2 x 3 min and submerged in 100%, 90%, and 70% EtOH for 5 min each. Slides were rinsed in dH2O and then submerged in Harris Haematoxylin (Sigma-Aldrich) for 3 min and rinsed in dH2O. Slides were then submerged in 95% EtOH for 30 seconds and dipped in Eosin Y (Abcam, UK) for 1 second. Stained sections were transferred to dH2O and then dehydrated in 70%, 90%, and 100% EtOH for 5 min each, followed by 5 min in 50% EtOH/xylene and then submerged twice in xylene for 3 min. DPX mounting media (Thermo Scientific™) was applied, cover slipped was applied and left to dry.

#### Assessment of intestinal crypt survival

Images were captured on a PANNORAMIC 250 Flash III DX slide scanner (3DHISTECH) as depicted in Fig 2.3. The CaseViewer (3DHISTECH, Version 2.4.0.119028) software was used to capture images at 5x and 20x magnification of mice FFPE small intestine and colon samples at all time points post-irradiation. The number of surviving crypts per circumference of the transverse sectioned intestine was enumerated from H&E-stained sections.

**SUPPLEMENTARY FIGURES**

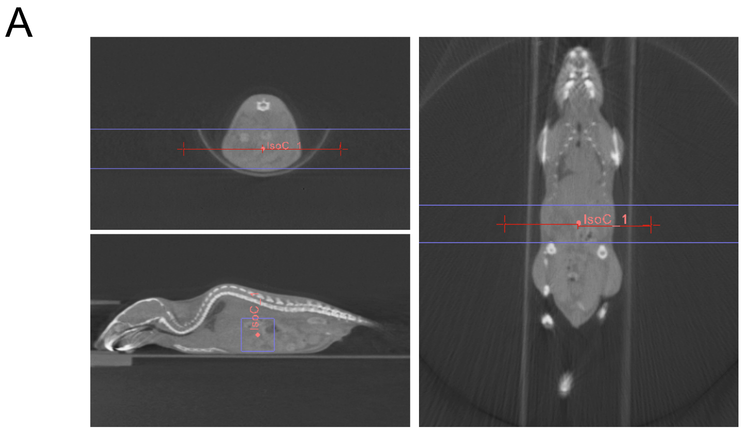


**Fig. S1 CT scan image to depict targeted region for X-ray delivery.**

CT scan guided single fraction dose of X-rays delivered within a 10 x 10 mm radiation field as depicted by the blue lines, with the isocentre depicted by the red dot. Care was taken to avoid the bone marrow and major organs with the CT scan images revealing an approximate 90-95% protection of the bone marrow by 100% protection of the spine and majority of the long bones.

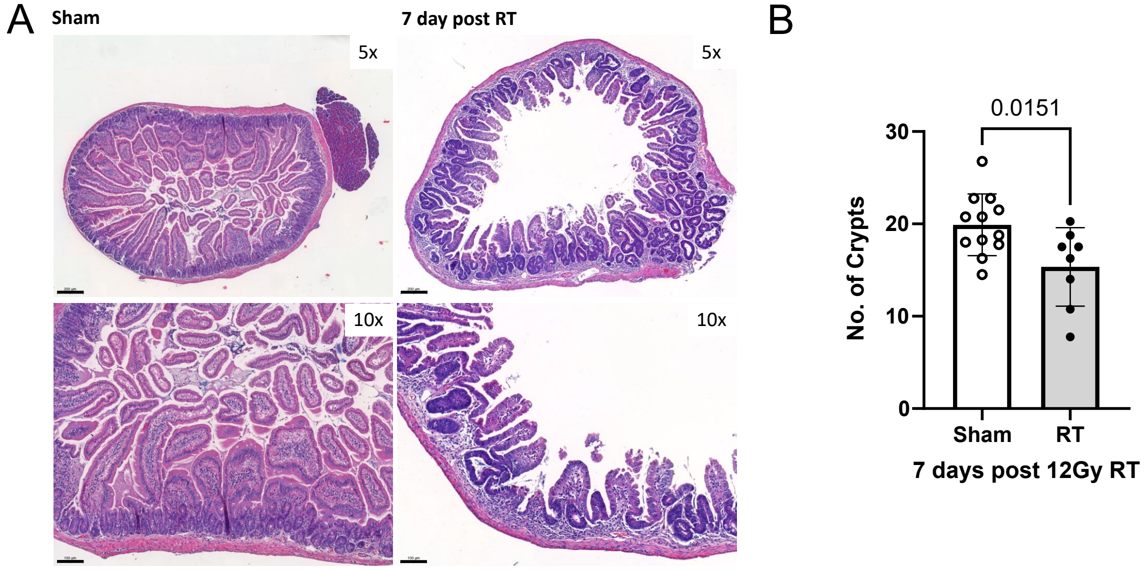


**Fig. S2 Localised abdominal RT results in intestinal crypt loss.**

(A) Representative images of the duodenum at day 7 post 12 Gy RT compared to sham at 5x and 10x magnification. Scale bar at 5x objective, 200 µm; 10x objective, 100 µm. (B) The average number of mean surviving crypts in the duodenum was quantified at 10x magnification view at day 7 post-RT, compared to sham mice. Data combined from more than 3 independent experiments n = 12 sham, and n = 8 RT. Significance quantified using unpaired parametric student's t-test.

29

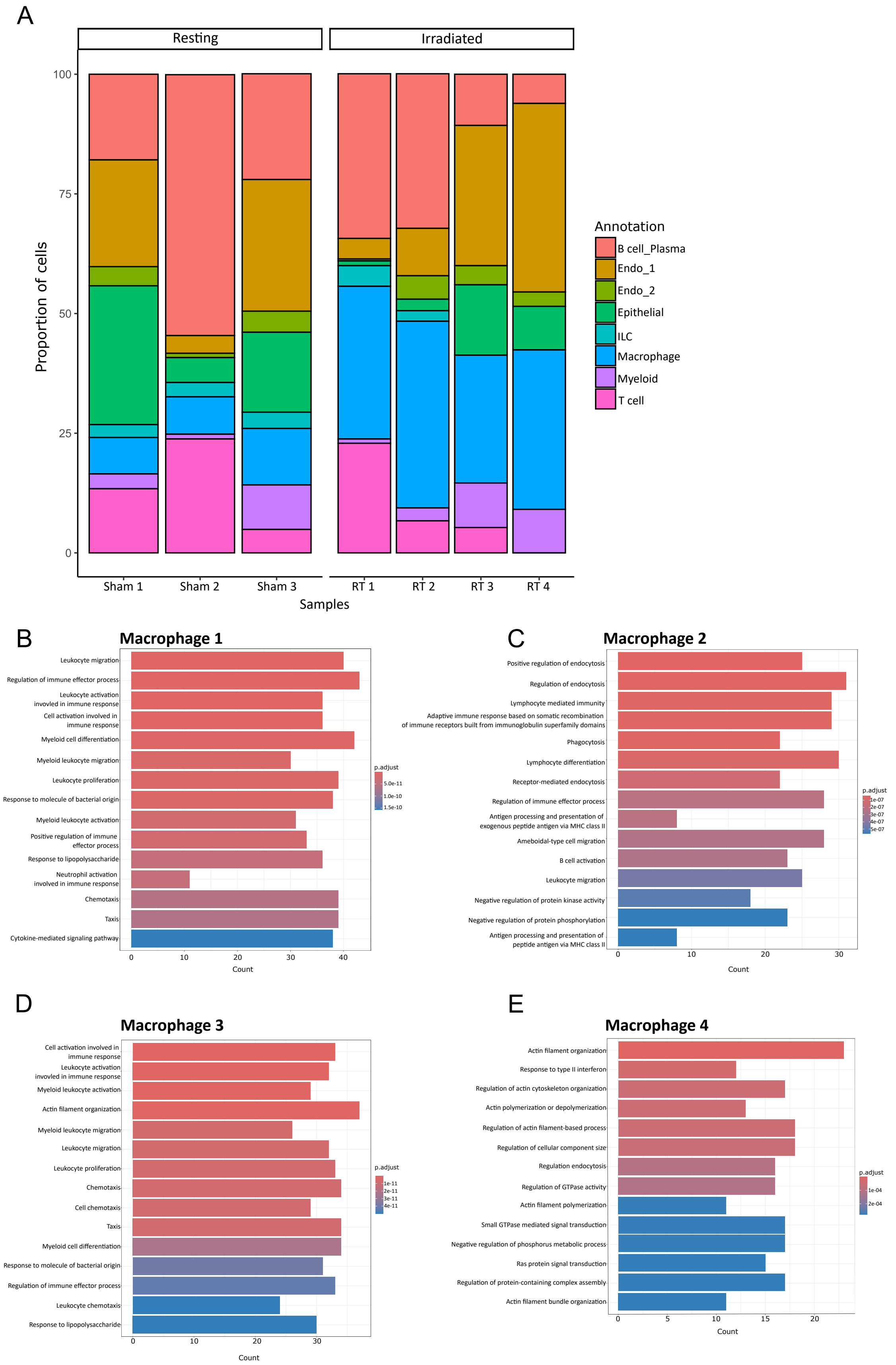
A
Resting
Irradiated
Proportion of cells
100
75
50
25
0
Sham 1
Sham 2
Sham 3
RT 1
RT 2
RT 3
RT 4
Samples
Annotation
B cell_Plasma
Endo_1
Endo_2
Epithelial
ILC
Macrophage
Myeloid
T cell
B
Macrophage 1
Leukocyte migration
Regulation of immune effector process
Leukocyte activation involved in immune response
Cell activation involved in immune response
Myeloid cell differentiation
Myeloid leukocyte migration
Leukocyte proliferation
Response to molecule of bacterial origin
Myeloid leukocyte activation
Positive regulation of immune effector process
Response to lipopolysaccharide
Neutrophil activation involved in immune response
Chemotaxis
Taxis
Cytokine-mediated signaling pathway
Count
p.adjust
C
Macrophage 2
Positive regulation of endocytosis
Regulation of endocytosis
Lymphocyte mediated immunity
Adaptive immune response based on somatic recombination of immune receptors built from immunoglobulin superfamily domains
Phagocytosis
Lymphocyte differentiation
Receptor-mediated endocytosis
Regulation of immune effector process
Antigen processing and presentation of exogenous peptide antigen via MHC class II
Ameboidal-type cell migration
B cell activation
Leukocyte migration
Negative regulation of protein kinase activity
Negative regulation of protein phosphorylation
Antigen processing and presentation of peptide antigen via MHC class II
Count
p.adjust
D
Macrophage 3
Cell activation involved in immune response
Leukocyte activation involved in immune response
Myeloid leukocyte activation
Actin filament organization
Myeloid leukocyte migration
Leukocyte migration
Leukocyte proliferation
Chemotaxis
Cell chemotaxis
Taxis
Myeloid cell differentiation
Response to molecule of bacterial origin
Regulation of immune effector process
Leukocyte chemotaxis
Response to lipopolysaccharide
Count
p.adjust
E
Macrophage 4
Actin filament organization
Response to type II interferon
Regulation of actin cytoskeleton organization
Actin polymerization or depolymerization
Regulation of actin filament-based process
Regulation of cellular component size
Regulation endocytosis
Regulation of GTPase activity
Actin filament polymerization
Small GTPase mediated signal transduction
Negative regulation of phosphorus metabolic process
Ras protein signal transduction
Regulation of protein-containing complex assembly
Actin filament bundle organization
Count
p.adjust

**Fig. S3 Irradiation-driven changes in the proportion of cell clusters observed through scRNA-seq.**

(A) Stacked bar chart shows the proportions of each cell subset in resting (n = 3) and irradiated small intestine (n = 4) samples at day 5 post-RT. Data includes n = 3 resting and n = 4 irradiated mice. Over-representation analysis of GO terms associated with (B) macrophage 1, (C) macrophage 2, (D) macrophage 3, and (E) macrophage 4 clusters.

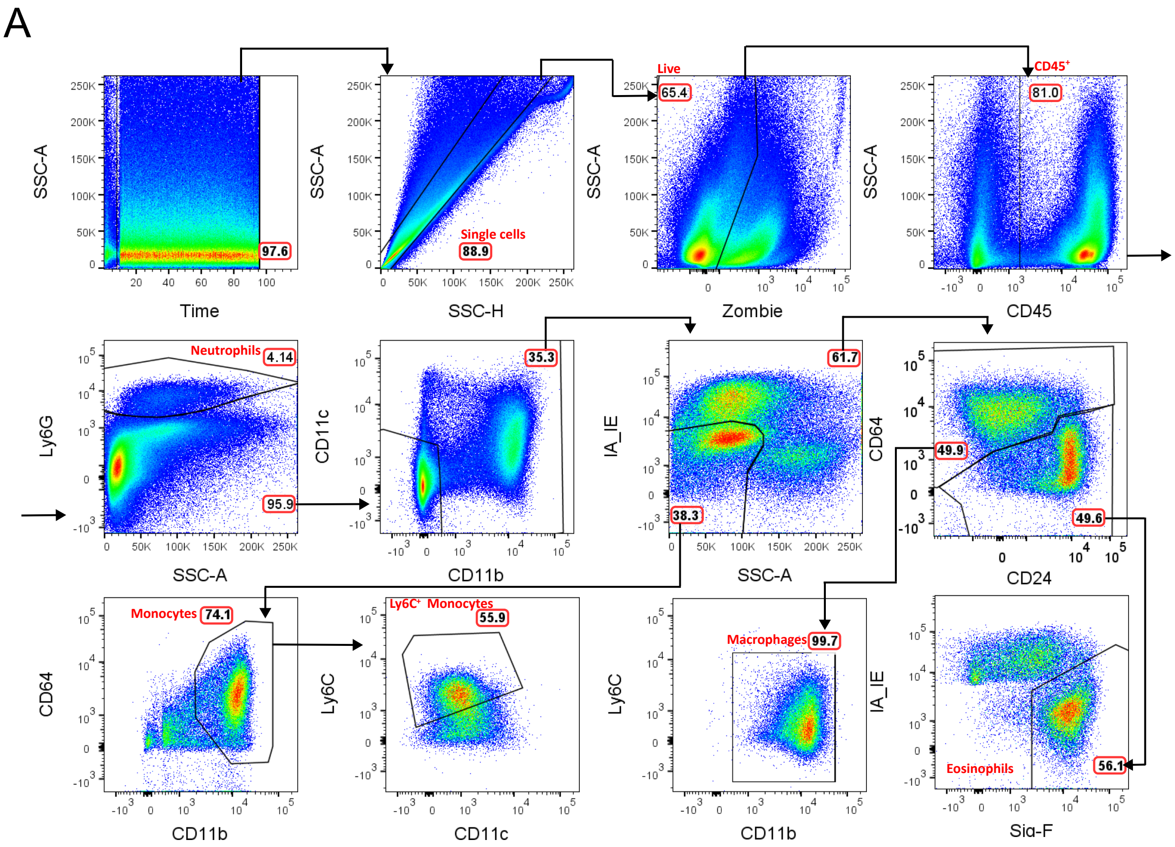


**Fig. S4 Flow cytometric gating strategy for identifying immune cell populations within the small intestine.**

(A) Single cells were isolated from the small intestine and stained with antibodies against cell surface markers. Immune cell populations were sequentially defined based on the expression profiles of the surface markers stated.

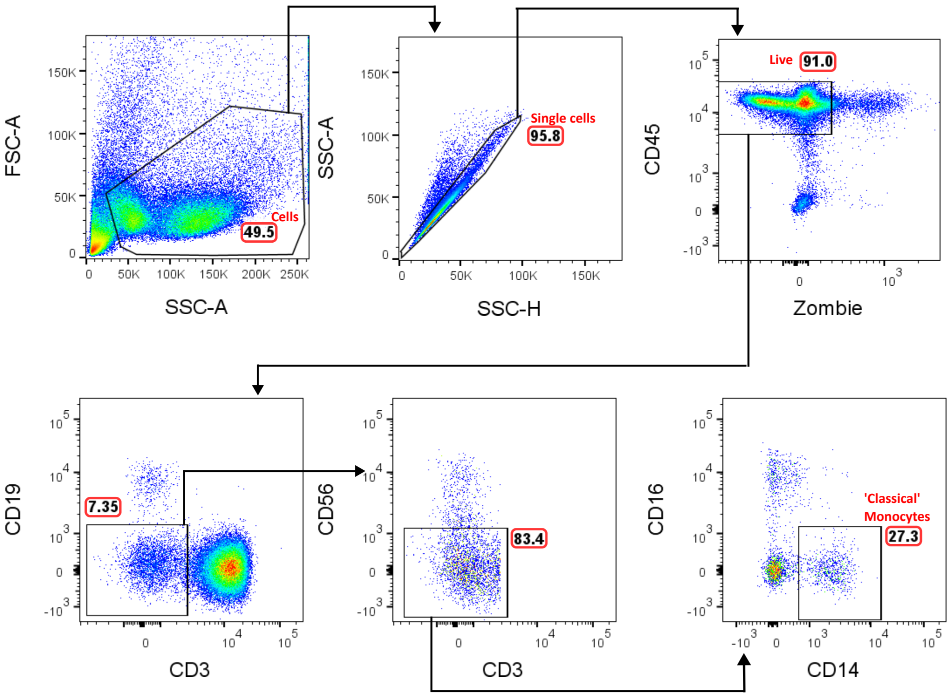


**Fig. S5 Flow cytometric gating strategy for identifying monocytes within the human blood.** Single cells were isolated from human PBMCs and stained with antibodies against human cell surface markers. 'Classical' monocytes were defined based on CD14 and CD16 expression and gated as $CD14^{+}CD16^{-}$.

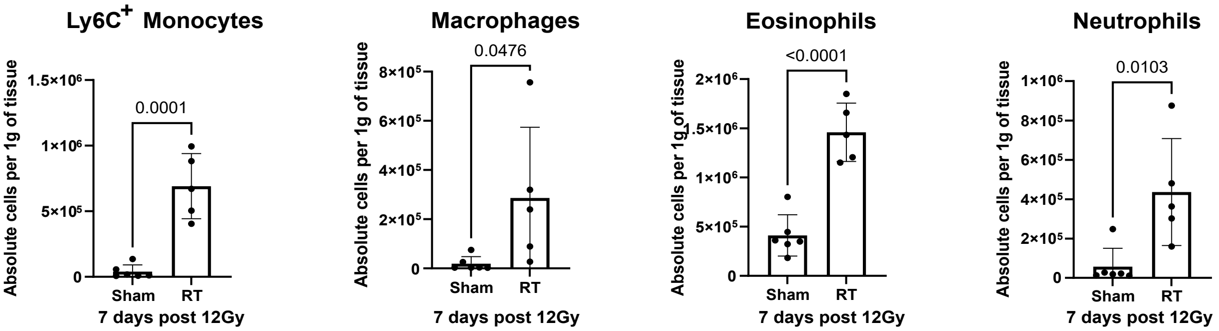


**Fig. S6 RT induces increased innate immune cell recruitment in iCCR reporter mice.** Flow-cytometric quantification $Ly6C^+$ monocytes, macrophages, eosinophils and neutrophils in sham and irradiated iCCR reporters at day 7 post-RT. Data was combined from 2 independent experiments, n = 6 sham and n = 5 RT iCCR reporters and analysed using unpaired parametric student's t-test. Each data point represents a measurement from a single mouse, and all numerical data represent mean (±SD).

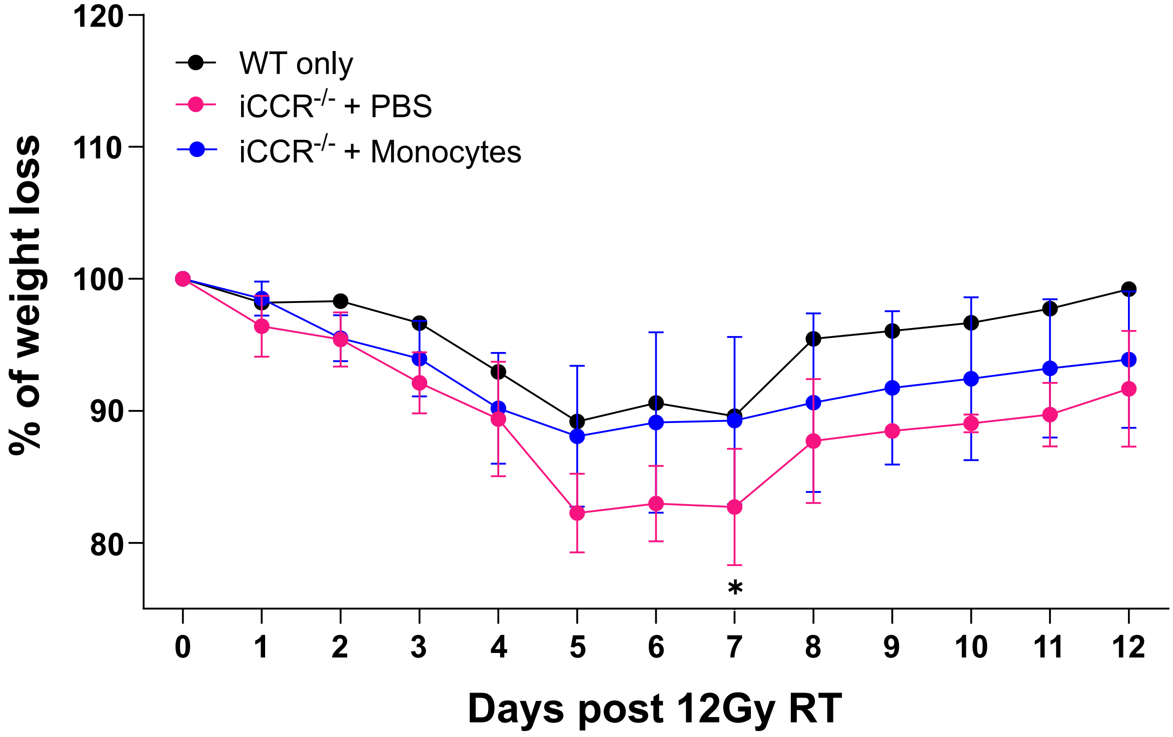


**Fig. S7 Adoptive transfer of Ly6C$^+$ monocytes is sufficient to protect against iCCR-dependent increased susceptibility to RT-induced weight loss.**

(A) Adoptive monocyte transfer of 1x10$^6$ monocytes into irradiated iCCR$^{-/-}$ at day 2 post-RT. Change in body weight of 12 Gy irradiated iCCR$^{-/-}$ mice injected with PBS and iCCR$^{-/-}$ injected with 1x10$^6$ Ly6C$^+$ monocytes, weights were recorded daily for up to 12 days. Statistical analysis using a two-way ANOVA with mixed-effects analysis, *$P < 0.05$. Data combined from more than 2 independent experiments, n = 12 irradiated WT, n = 16 irradiated iCCR$^{-/-}$ injected with PBS, n = 4 irradiated iCCR$^{-/-}$ injected with 1x10$^6$ Ly6C$^+$ monocytes. Each data point represents a measurement from a single mouse, and all numerical data represent mean (±SD).

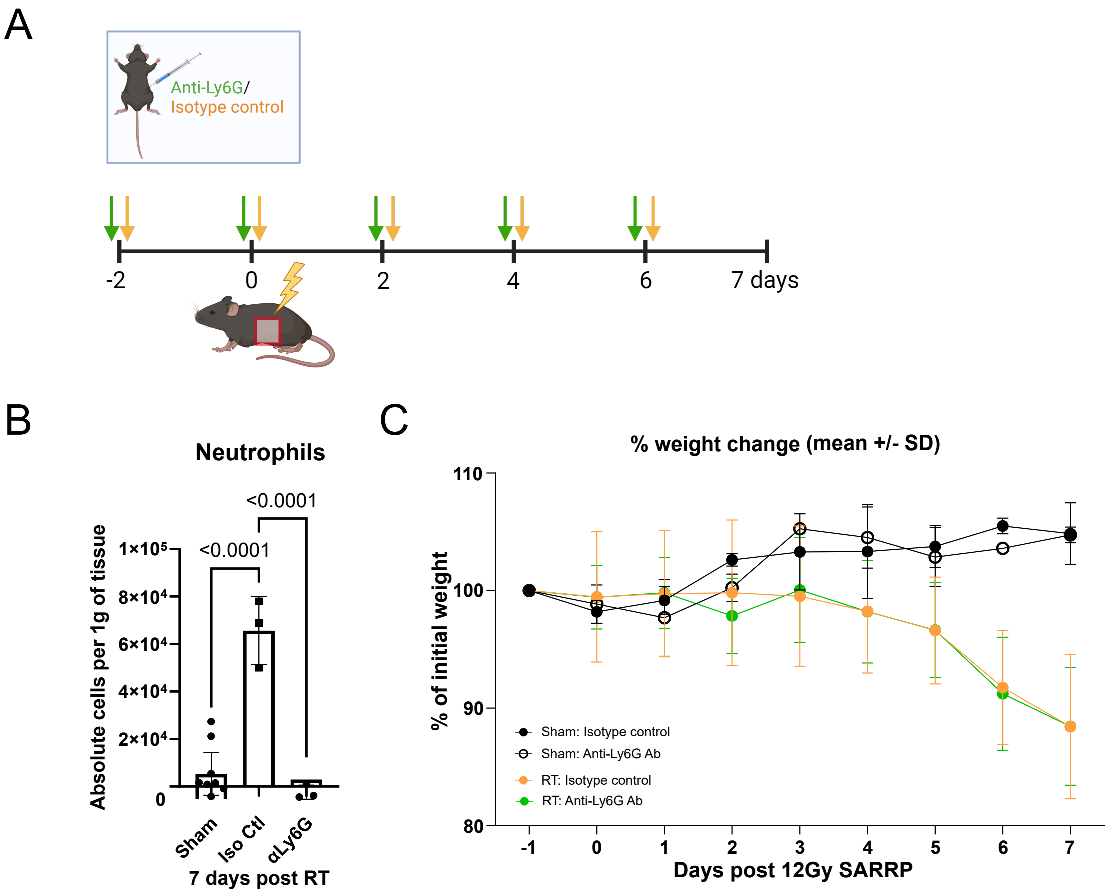


**Fig. S8 The inability to recruit neutrophils into the small intestine does not affect the extent of RT-induced weight loss.**

(A) I.P. injections of either isotype control or anti-Ly6G mAb started two days before single fraction 12 Gy RT and then every two days on alternating sides of the mouse abdomen. (B) Flow cytometric quantification of neutrophils in the small intestine in sham and irradiated isotype control mice and sham and irradiated anti-Ly6G treated mice at day 7 post-RT. Statistical analysis using ordinary one-way ANOVA with Tukey's multiple comparisons test. Each data point represents a measurement from a single mouse, and all numerical data represent mean (±SD). (C) Change in body weight of 12 Gy irradiated C57BL/6 mice treated with either isotype control or anti-Ly6G mAb. Weights were recorded daily for up to 7 days. Statistical analysis using two-way ANOVA with mixed-effects analysis. Data from one experiment, n = 2 sham + isotype control, n 2 = sham + anti-Ly6G mAb, n = 6 RT + isotype control, n = 6 RT + anti-Ly6G mAb.

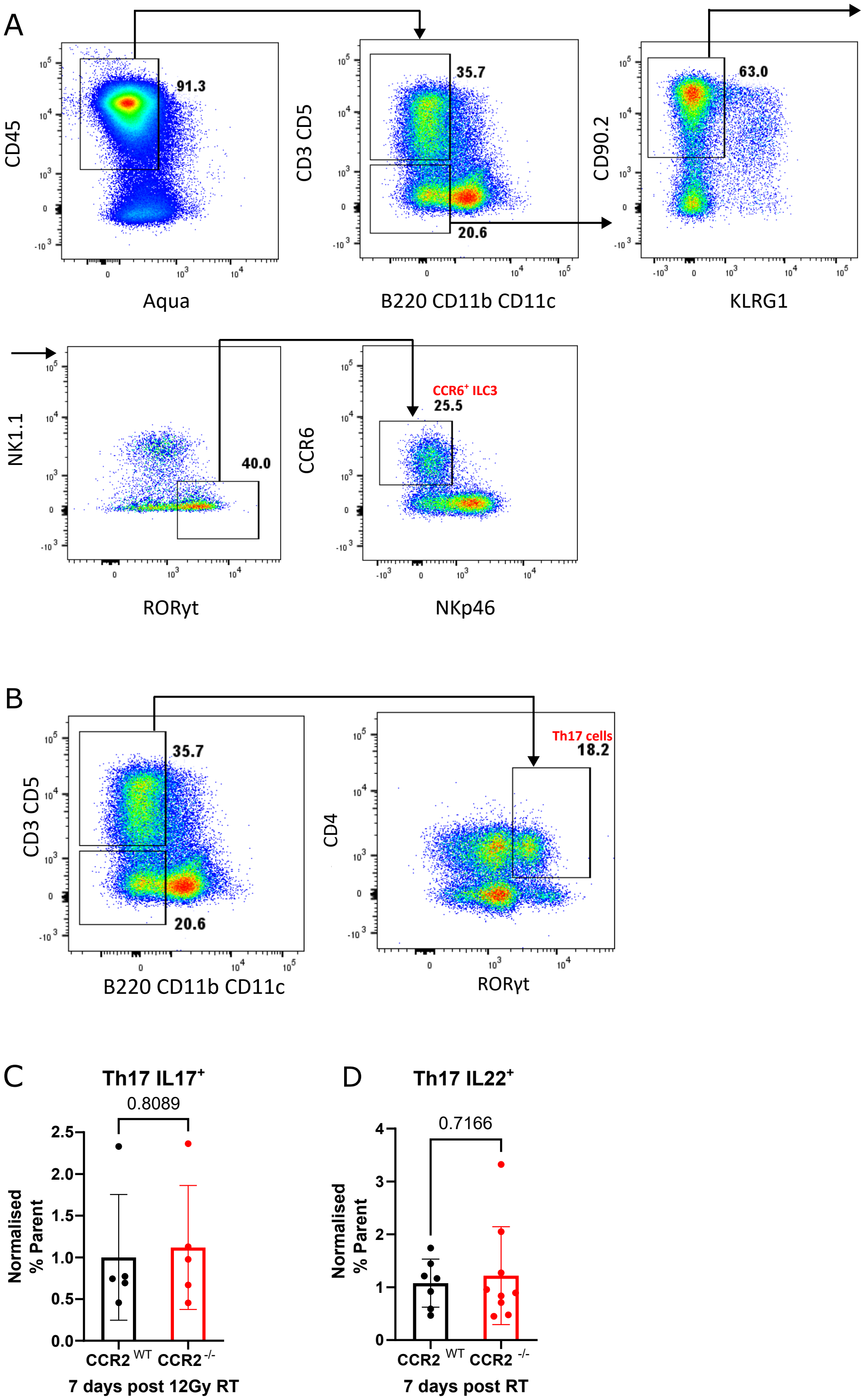
A
CD45
Aqua
91.3
CD3 CD5
B220 CD11b CD11c
35.7
20.6
CD90.2
KLRG1
63.0
NK1.1
RORγt
40.0
CCR6
NKp46
CCR6+ ILC3
25.5
B
CD3 CD5
B220 CD11b CD11c
35.7
20.6
CD4
RORγt
Th17 cells
18.2
C
Th17 IL17+
0.8089
Normalised % Parent
CCR2 WT
CCR2 -/-
7 days post 12Gy RT
D
Th17 IL22+
0.7166
Normalised % Parent
CCR2 WT
CCR2 -/-
7 days post RT

73

**Fig. S9 Production of IL-17 and IL-22 from intestinal ILC3 and Th17 cells in CCR2$^{-/-}$ and WT mice at day 7 post-RT.**

(A) Flow cytometry gating strategy for intestinal ILC3 populations expressing IL-17 and IL-22. (B) Flow cytometry gating strategy for IL-17A and IL-22 expression on RORyt$^{+}$CD4$^{+}$ Th17 cells. Normalised frequency of Th17 cells expressing (C) IL-17A and (D) IL-22. All graphs compare WT and CCR2$^{-/-}$ mice at 7 days post 12 Gy RT. Data combined from more than 2 independent experiments and includes at least n = 5 in each group. Data was analysed with an unpaired nonparametric t-test using Mann Whitney test. Each data point represents a measurement from a single mouse, and all numerical data represent mean (±SD).
